# NDUFA4L2 rescues hyperoxia-induced migration defects in retinal endothelial cells by reversing isocitrate dehydrogenase flux blockade

**DOI:** 10.64898/2026.07.14.738274

**Authors:** Hyungryun Jang, Ashwin Chandra, Kobe Tray, Brooke Linnehan, Fabian Schulte, Gopalan Gnanaguru, Charandeep Singh

## Abstract

Retinopathy of prematurity (ROP) is caused by hyperoxic exposure of prematurely born infants. The mouse model of oxygen-induced retinopathy (OIR) recapitulates pathological features of both phase I and phase II ROP. We here looked at the retinal proteins that change in response to hyperoxia in phase I of the mouse model of OIR. Using tandem mass tag labeled proteomics, we found several differentially expressed proteins (DEPs) in phase I of OIR. Of all the DEPs, we investigated the role of previously unknown protein NADH dehydrogenase [ubiquinone] 1 alpha subcomplex subunit 4-like 2 (NDUFA4L2). NDUFA4L2 protein and its paralog NDUFA4 are both mitochondrial complex I proteins; however, here we demonstrate that NDUFA4L2 changes in both phases of OIR, with no changes in its paralog NDUFA4, implying its unique function in pathophysiology of the disease. We demonstrate that NDUFA4L2 is an oxygen-sensitive protein and regulates retinal endothelial cell migration by rescuing isocitrate dehydrogenase flux impaired by hyperoxia in phase I of OIR.

## Introduction

Supplemental oxygen is administered to high-risk prematurely born infants until they are sufficiently able to keep up their PO2 levels[1–3]. Although this supplemental oxygen or hyperoxia is necessary to prevent mortality in infants who have insufficiently developed lungs, it leads to downregulation of neurovascular development in growing premature infants, leading to vaso-obliteration in the retina, causing phase I of retinopathy of prematurity (ROP) [3–5]. When these infants are sufficiently able to maintain their PO2 levels, these infants are relieved of supplemental oxygen, and this pseudohypoxia after moving from hyperoxic to normoxic conditions leads to the neovascularization in the retina, causing phase II of ROP [3–5]. As per 2010 estimates 184,700 infants were born premature, with 32,300 developing ROP [6]. ROP incidences in the US alone have doubled from 2003-2019 [7]. These numbers add up yearly and as per 2010 a total of 19 million children were visually impaired, including those suffering from diseases other than ROP [8]. Current ROP treatments are limited to treating the second phase of ROP, either by destroying the neovascular region using laser photocoagulation or by inhibiting neovascular pathological angiogenesis using anti-vascular endothelial growth factor (VEGF) antibodies [5, 9]. Anti-VEGF treatment is widely used to treat ROP in phase II because it effectively prevents further exacerbation of the vascular damage. However, excessive anti-VEGF treatment can hinder normal vascular development and vascular integrity, and anti-VEGF treatment requires long follow ups to keep track of any reactivation of vascular abnormalities in later timepoints [10]. We and others have demonstrated that phase I is the precursor to phase II of ROP and oxygen-induced retinopathy (OIR) [11–14]. Preventing ROP in Phase I can prevent vision loss before it worsens to phase II, to an extent where a cure merely means saving the remaining intact vision rather than reversing the phenotype. There is no treatment available yet to prevent ROP early in phase I. It has been demonstrated in the mouse model of OIR that hyperoxia primarily impacts the developing vasculature in phase I by inhibiting endothelial cell proliferation and migration [15, 16]. Endothelial cell (EC) migration and proliferation are both energy-demanding metabolic processes. ECs generate close to 85% of energy from glucose [17]. Inhibition of 6-phosphofructo-2-kinase/fructose-2,6-bisphosphatase 3 (PFKFB3), a glycolysis regulator, stalls vascular development [17]. Modulating PFKFB3 levels modulates angiogenesis in a mitogen-independent manner [18, 19], demonstrating that glycolytic regulation is sufficient to regulate angiogenesis. Broadly speaking, glycolysis controls migration and glutamine metabolism via tricarboxylic acid (TCA) cycle controls proliferation of endothelial cells. However, alpha-ketoglutarate (αKG) in TCA cycle is an exception to this broadly accepted glycolysis vs. TCA contributions, with αKG contributing to both proliferation and migration [20, 21]. Moreover, low birth weight is associated with the worst ROP outcomes in prematurely born infants, implying that infants who face metabolic challenges are more susceptible to ROP [22].

Vascular development requires a hypoxic environment, which leads to the activation of VEGF-VEGF receptor (VEGFR) signaling pathway, which is the primary regulator of angiogenesis. VEGF is produced by astrocytes, retinal ganglion cells, and, to a small extent, by Müller cells and drives vascular patterning, proliferation, and migration of endothelial cells in the retina [23, 24]. Of all the known VEGF isoforms, VEGFA dictates retinal angiogenesis by binding to VEGFR1 and VEGFR2 on the endothelial cell surface [25, 26]. VEGFR2 is required for tip cell selection, migration, and proliferation of endothelial cells, whereas VEGFR1 fine-tunes these processes [25, 26]. Soluble VEGFR1 secreted by stalk ECs helps neutralize VEGFA in proximity, to further fine-tune the VEGF gradient and guide vascular sprouting by negatively regulating the conversion of stalk cells into tip cells [27]. VEGF signaling is not limited to ECs. VEGF secreted by nerves controls neurovascular patterning by acting on vascular neuropilin 1 (Nrp1) during development [28]. VEGF also mediates axonal guidance in response to VEGFR2 present on neuronal granule cells and modulates neuronal N-methyl-D-aspartate (NMDA) activity [26]. Although VEGF production goes down in hyperoxia and is blamed for being a culprit in hyperoxia-mediated vaso-obliteration, supplementing VEGF and other growth factors like epidermal growth factor (EGF) and insulin-like growth factor (IGF), in culture doesn’t rescue vascular proliferation and migration [29, 30], implying the mechanistic underpinning in OIR is not yet fully understood. The problem can be considered two folds, first is the downregulation of mitogens like VEGF, and second is the aberrant perception of mitogens by their corresponding receptors like VEGFR.

In this study, we aimed to understand the problem in the perception of VEGF signaling by ECs, particularly in an attempt to understand the mechanism underlying the phase I pathophysiology. We used both the in vivo and in vitro models and found a previously unknown protein, NADH dehydrogenase [ubiquinone] 1 alpha subcomplex subunit 4-like 2 (NDUFA4L2) which is downregulated in hyperoxic phase I and upregulated in hypoxic phase II of OIR. NDUFA4L2 is known to impact proliferation and act by modulating metabolism in cancers [31–34]; however, its physiological function in endothelial cells and ROP is not known. We here demonstrate the dual metabolic function of NDUFA4L2, and the mechanistic control of migration by modulating isocitrate dehydrogenase (IDH) flux.

## Results

### Tandem Mass Tag (TMT)-labeled proteomic analysis of mouse retina showed downregulation of a previously unknown mitochondrial protein target in phase 1 of OIR

To find out proteomic changes induced in response to hyperoxia, we performed TMT-labeled proteomics analysis of P12 phase I OIR-modeled mouse retina. Mice were exposed to normoxic room air or hyperoxic 75% oxygen tension from postnatal day (P)7-P12. Mice were euthanized on P12, eyes were enucleated, and retinas were dissected out of the eyes under the microscope, while maintaining samples on ice. Proteins were extracted from the retina, TMT-labeled, fractionated, and then analyzed on a high-resolution mass spectrometer. A total of 7704 proteins were identified in the samples. A total of 19 proteins were downregulated in hyperoxic samples with a fold change of 1.5 and a p-value of 0.05. Whereas a total of 522 proteins were upregulated in hyperoxic samples with a fold change of 1.5 and a p-value of 0.05. These changes are depicted in the volcano plot in Figure 1a. All the statistically significantly changed differentially expressed proteins (DEPs) were parsed through MouseMine [35] to look at the pathway enrichment. Pathway analysis showed the upregulated DEPs were enriched in neuronal changes (Supplemental table1), which we speculate to be a consequence of vaso-obliteration in phase 1. Since there were only 19 downregulated DEPs, pathway enrichment didn’t yield any output. We checked the Uniprot annotation score for all the up- and down-regulated DEPs. Of all the DEPs, one of the previously uncharacterized proteins, NDUFA4L2, had the least UniProt score of 1/5, meaning its function is least understood. NDUFA4L2 was found to be downregulated in P12 hyperoxic retinas as compared to room air P12 retinas in the TMT-proteomics data (Figure 1a and b). Whereas its paralog NDUFA4 didn’t show a significant difference in our proteomics data set (Figure 1b). We also looked at the published P17 phase II OIR data set GSE234447 [36], and as expected based on our P12 findings, *NDUFA4L2* transcription levels showed a trend opposite to that seen in our phase I data (Figure 1c), whereas, similar to our dataset, the P17 dataset also showed no differences in *NDUFA4* transcription levels (Figure 1c).

**Figure 1.**
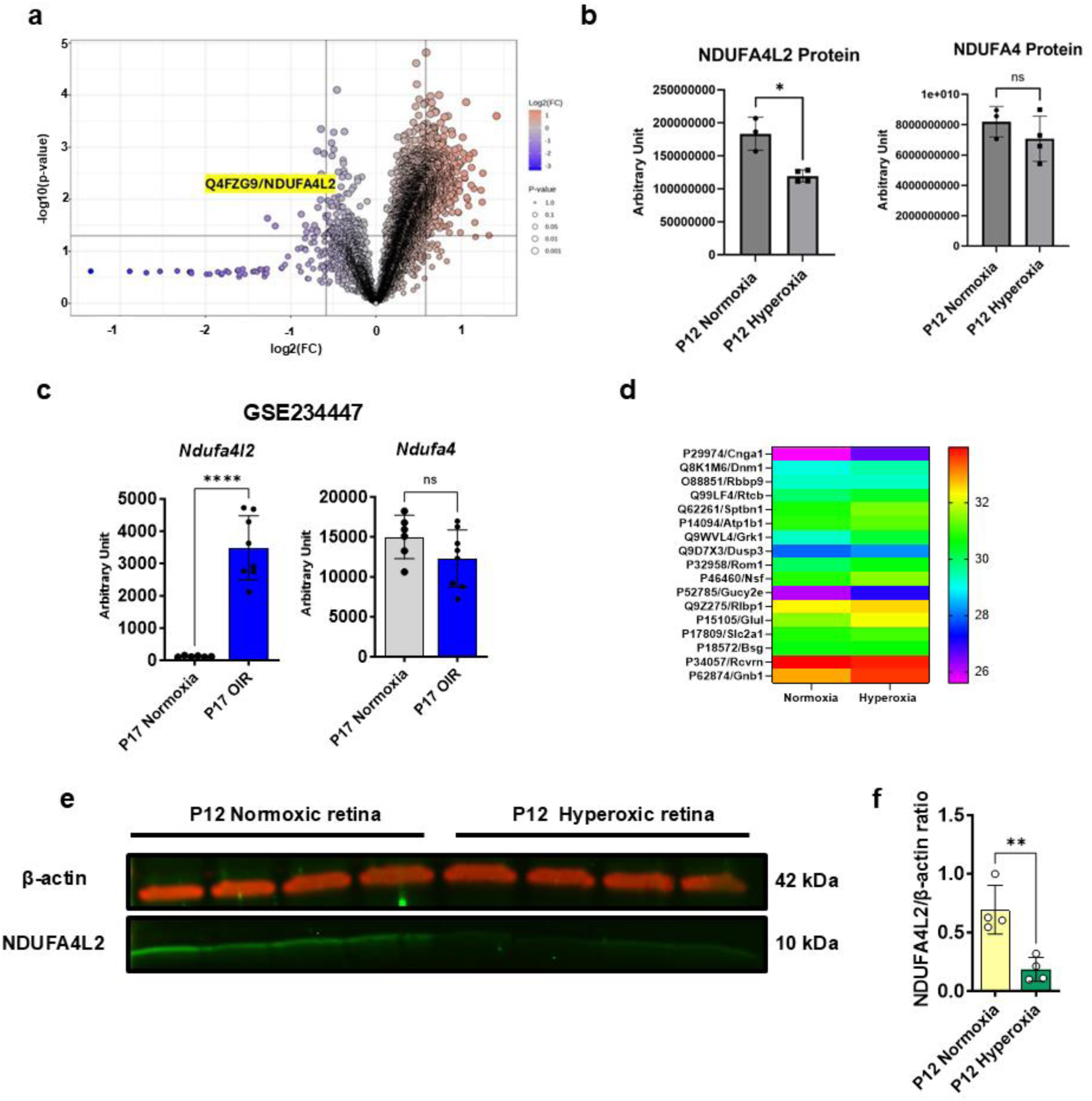
NDUFA4L2 is downregulated in phase I of oxygen-induced retinopathy (OIR) a) Volcano plot of TMT-labeled proteomics data showed downregulation of NDUFA4L2 in P12 hyperoxia OIR mouse retina compared to normoxic controls. b) TMT-labeled proteomics data demonstrated downregulation of NDUFA4L2 protein and no changes to its paralog NDUFA4 in P12 OIR mouse retinas in hyperoxic conditions. c) Re-analysis of GSE234447 data showed *Ndufa4l2* transcript levels in phase II OIR mouse retina (P17) demonstrated upregulation in response to pseudohypoxia in phase II, opposite to that observed in our phase I dataset, whereas *Ndufa4* transcript levels did not differ significantly in the P17 phase II OIR dataset. d) Our phase I OIR hyperoxic dataset demonstrated an opposite trend to multiple other targets seen in a published study pertaining to phase II OIR. e) and f) Western blotting of retinal lysates from a different P12 OIR mouse cohort confirmed our findings from the TMT-labeling experiment, which showed downregulation of NDUFA4L2 in P12 hyperoxic OIR retinas. Data are presented as mean ± SD (n = 4 per group). Statistical significance was determined using an unpaired two-tailed Welch’s t-test. Statistical significance is indicated as: *p-value ≤ 0.05, **p-value ≤ 0.01, and ****p-value ≤ 0.0001. NS indicates no significant difference.

To benchmark our TMT-labeled proteomics dataset, we compared a published proteomics dataset by Vähätupa et al.[37] to statistically significant changes in our proteomics dataset. Vähätupa et al.[37] studied proteomics changes in the retinal proteome of P13 OIR mice. Since P13 is one day post-hyperoxic exposure in the OIR model, which is an early phase of pseudohypoxia, we expected to see changes opposite to those seen in our hyperoxic P12 OIR proteomics data. We looked at the proteins found to be statistically significant and differentially expressed proteins in Vähätupa et al. in our data. The data is plotted in Figure 1d. All the proteins reported in the P13 DEP data set showed opposite and statistically significant differences, except retinoblastoma binding protein 9 (RBBP9), retinaldehyde binding protein 1 (RLBP1), basigin (BSG), and recoverin (RCVRN), which were not statistically significantly changed in our dataset, showing consistent changes in proteins in the two datasets.

### NDUFA4L2 is an oxygen-sensitive protein target in retinal endothelial cells

The role of NDUFA4L2 has been studied in many cancer types [34, 38–40]. However, its physiological role in the eukaryotic cells and its role in the pathophysiology of OIR have not been investigated so far. To confirm our findings from untargeted mass spec-based proteomics, we performed a secondary targeted measurement of NDUFA4L2 protein in P12 hyperoxic retinas and compared them with age-matched normoxic retinas, with the help of western blotting. Western blots showed downregulation of NDUFA4L2 in hyperoxic retina samples as compared to normoxic samples, indicating that the protein is sensitive to oxygen exposure, further confirming our TMT-labeled proteomics results (Figure 1e and f).

Since the OIR model is an angiogenesis model in which hyperoxic exposure leads to downregulation of angiogenesis, we speculated that NDUFA4L2 is an angiogenic target, as it is downregulated in the retina of hyperoxic OIR modeled mice. We checked the published mouse retinal single-cell RNA-sequencing dataset (GSE243413 [41]) and found that *Ndufa4l2* is predominantly expressed in retinal endothelial cells and pericytes (Figure 2a-c). We further performed an immunohistochemistry (IHC) antibody labeling experiment to evaluate whether NDUFA4L2 co-localizes with endothelial cells in the retina flat mounts. NDUFA4L2 appeared to co-localize with retinal blood vessels and showed punctuated labeling, indicating some low levels of expression in other cell types too or non-specific bindings of antibody (Supplemental Figure S1a and b). Interestingly, NDUFA4L2 showed an oxygen-dependent localization in retinal flat mounts, with a clear labeling in arteries but very little to no labeling in the vein in normoxic P12 mice retina (Supplemental Figure S1a and b). These results further established the endothelial-specific function of NDUFA4L2 and its role in the physiological oxygen gradient. Since retinal flat mount labeling can result from cells other than target cells if the cells are in proximity, we further looked at the expression of NDUFA4L2 in cultured primary human retinal endothelial cells (HRECs). Interestingly, we found very high basal expression of NDUFA4L2 in primary HRECs, and its downregulation in hyperoxic conditions (Figure 2d and e).

**Figure 2.**
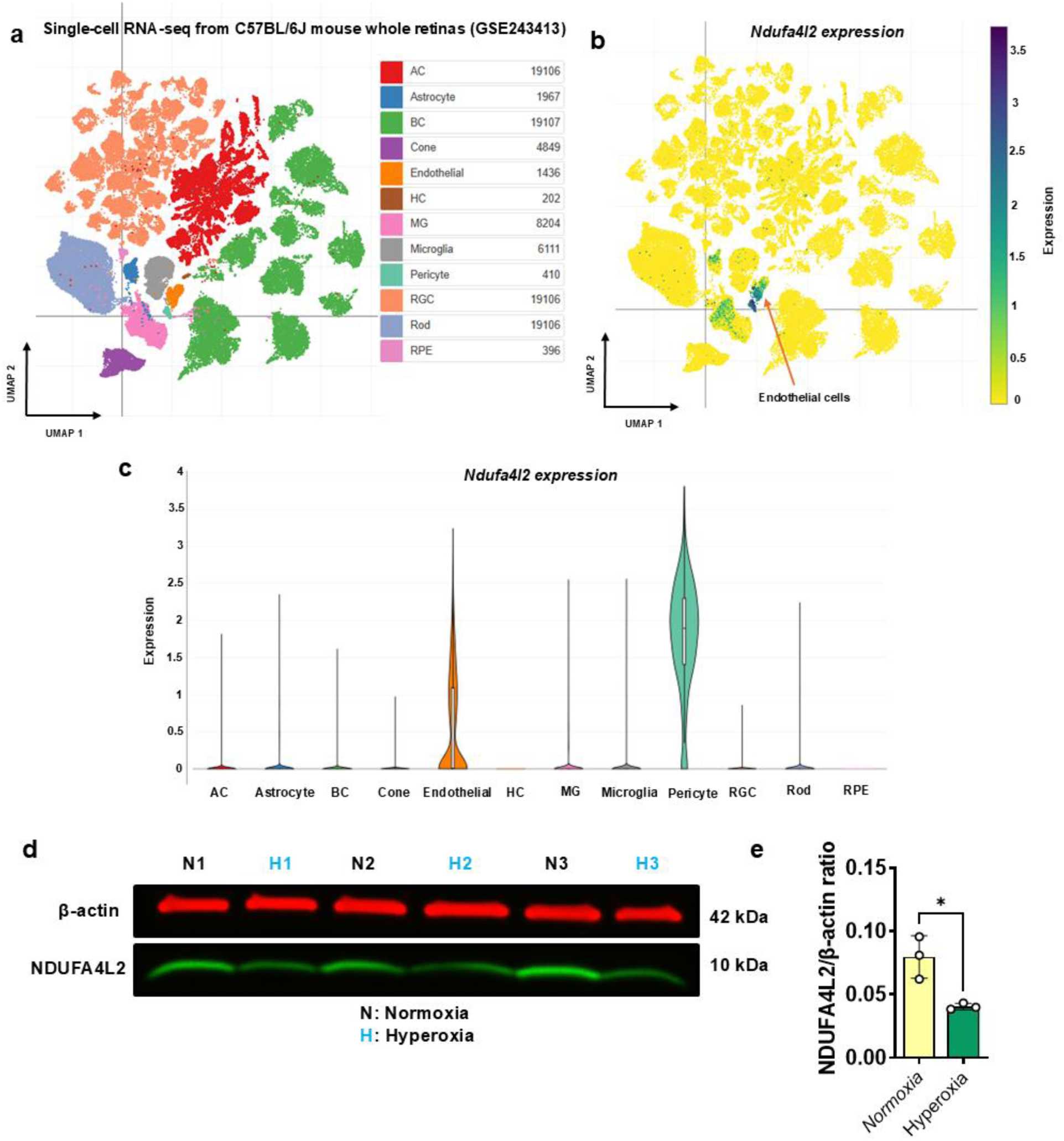
NDUFA4L2 expression is sensitive to hyperoxia in retinal endothelial cells. a-c) Re-analysis of the published dataset GSE243413 on Broad Institute Single Cell Portal [65] shows expression of *Ndufa4l2* in different retinal cell types, with the highest expression in endothelial cells and pericytes. d) Primary HRECs exposed to hyperoxia (H) for 48 h showed downregulation of NDUFA4L2 in comparison to normoxia (N), further confirming our findings that NDUFA4L2 is an endothelial-specific protein. e) quantification of NDUFA4L2 western blot image. Data are presented as mean ± SD (n = 3 per group). Statistical significance was determined using an unpaired two-tailed Welch’s t-test. Statistical significance is indicated as: *p-value ≤ 0.05.

### NDUFA4L2 is not under the control of hypoxia-inducible factor 1 alpha (HIF1α) at protein levels in retinal endothelial cells

Since hyperoxia downregulates HIF1α [42, 43] and has been shown to be upstream of NDUFA4L2 in cancers [34, 38–40], we hypothesized that NDUFA4L2 may be a HIF1α target in endothelial cells, too. To test if NDUFA4L2 is an HIF1α target, we used two different pharmacological compounds to stabilize HIF1α. We first used MG132, which upregulates HIF1α by inhibiting proteasomal degradation of ubiquitinated HIF1α [44, 45]. The low dose of MG132 treatment of primary HRECs led to the stabilization of HIF1α protein in normoxic conditions, as expected (Figure 3a and b). Interestingly, at protein levels, MG132 didn’t increase NDUFA4L2 levels when HIF1α was upregulated, implying NDUFA4L2 protein is regulated in a HIF1α-independent manner (Figure 3c and d). The low dose of MG132 we used was able to upregulate transcript levels of *NDUFA4L2* in normoxic conditions, however, it failed to upregulate HIF1α protein and *NDUFA4L2* transcript in hyperoxic conditions (Supplemental Figure S2a). This implies that the HIF1α only controls NDUFA4L2 transcription, not the protein, in normoxic conditions but not in hyperoxia. Although MG132 is a proteasomal inhibitor and HIF1α stabilization is indirect, we still looked at HIF1α target genes to make sure HIF1α was active in MG132-treated cells. We were able to see upregulation of signatures of HIF1α activity in MG132- treated cells (Supplemental Figure S2a).

**Figure 3.**
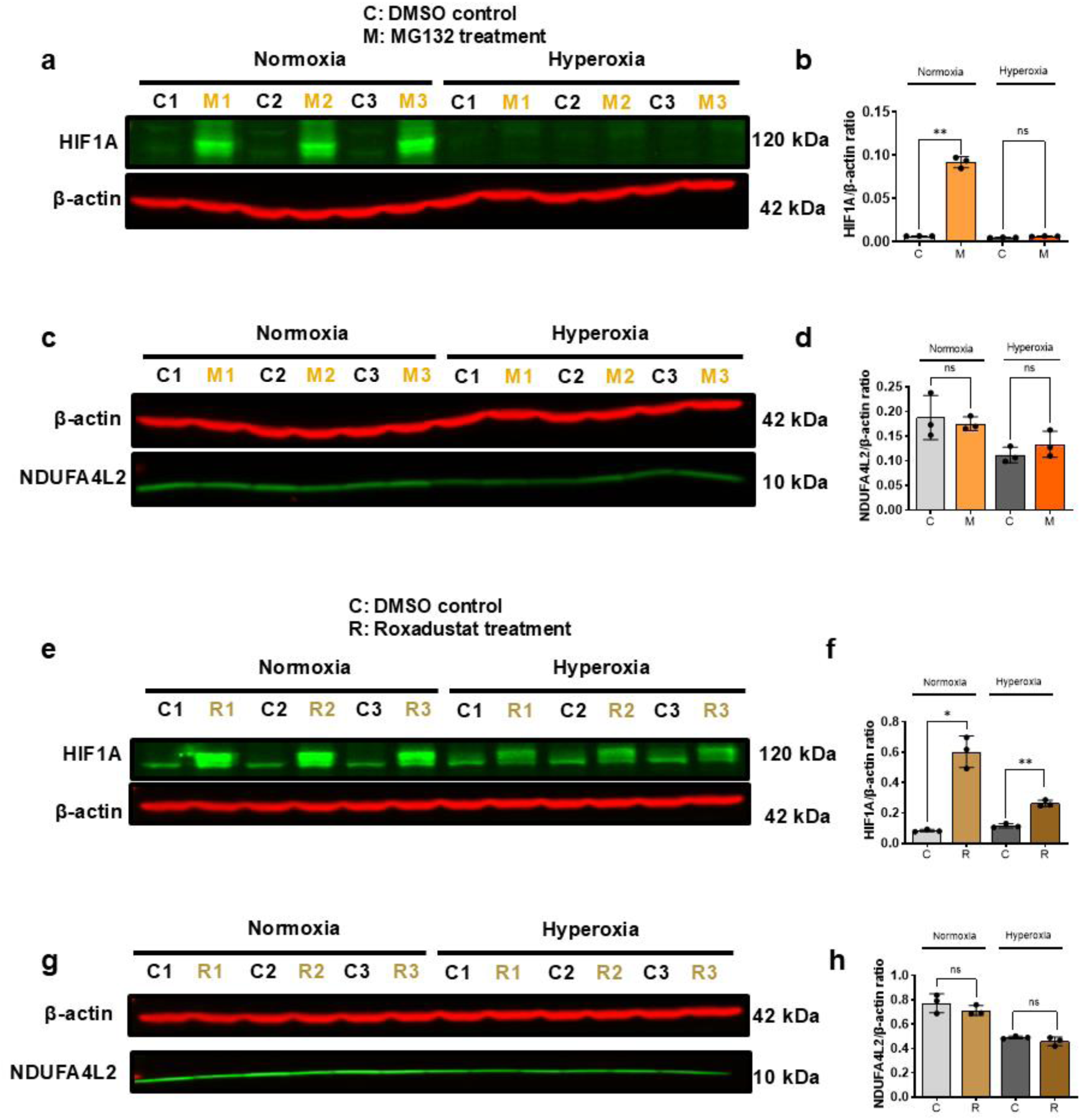
NDUFA4L2 is not the direct downstream target of HIF1α. a) The low dose of MG132-treated HRECs showed higher expression of HIF1α in normoxia, as expected. b) quantification of HIF1α western blot with or without MG132 treatment showing differences in HIF1α in response to MG132. c) NDUFA4L2 didn’t show a statistically significant increase in MG132-treated primary HRECs. d) quantification of NDUFA4L2 western blot with or without MG132. e) Roxadustat-treated HRECs showed higher expression of HIF1α as expected. f) quantification of HIF1α western blot with or without Roxadustat treatment g) NDUFA4L2 did not show a statistically significant change in Roxadustat-treated cells. h) quantification of NDUFA4L2 western blot with or without Roxadustat treatment. Data are presented as mean ± SD (n = 3 per group). Statistical significance was determined using an unpaired two-tailed Welch’s t-test. Statistical significance is indicated as: *p-value ≤ 0.05 and **p-value ≤ 0.01. NS indicates no significant difference. Legends: C, DMSO control; M, MG132 treatment; R, Roxadustat treatment.

Since MG132 is not a very specific HIF1α stabilizer and can prevent degradation of other proteasomal targets, we used a secondary method to confirm our findings. We used a very potent prolyl hydroxylase (PHD) inhibitor, Roxadustat, which stabilizes HIF1α by inhibiting its hydroxylation by PHD [12, 46]. Roxadustat binds to PHD’s active domain and prevents PHD-dependent hydroxylation of HIF1α, downstream ubiquitination, and proteasomal degradation. Western blot analysis of the proteins demonstrated stabilization of HIF1α in Roxadustat-treated cells (Figure 3e and f) and showed upregulation of HIF1α targets (Supplemental Figure S2b). However, NDUFA4L2 protein levels didn’t correlate with the HIF1α levels, implying NDUFA4L2 is not a HIF1α target in primary HRECs (Figure 3g and h).

### NDUFA4L2 protein levels are regulated independently of direct VEGFR2 signaling despite VEGF-driven transcriptional induction

Since we could not find HIF1α-dependent regulation of NDUFA4L2 in primary HRECs, we speculated that this could be a target of a downstream protein. One of the major players in endothelial hypoxia signaling and vascular development, other than the HIF1α, is VEGF. VEGFR1 and VEGFR2 are two main receptors present on the retinal endothelial cells, with VEGFR2 being the most important regulator of proliferation and migration [47]. We first tested whether hyperoxia impacts VEGFR2 levels in primary HRECs. Immunocytochemistry confirmed that hyperoxia significantly reduced VEGFR2 levels compared to normoxia (Figure 4a and b). To understand if VEGFR2 is upstream of NDUFA4L2, we treated primary HRECs with Apatinib, a selective tyrosine kinase inhibitor of VEGFR2. After 72 hours, 20 µM of Apatinib led to the downregulation of NDUFA4L2 protein levels (Figure 4c and d).

**Figure 4.**
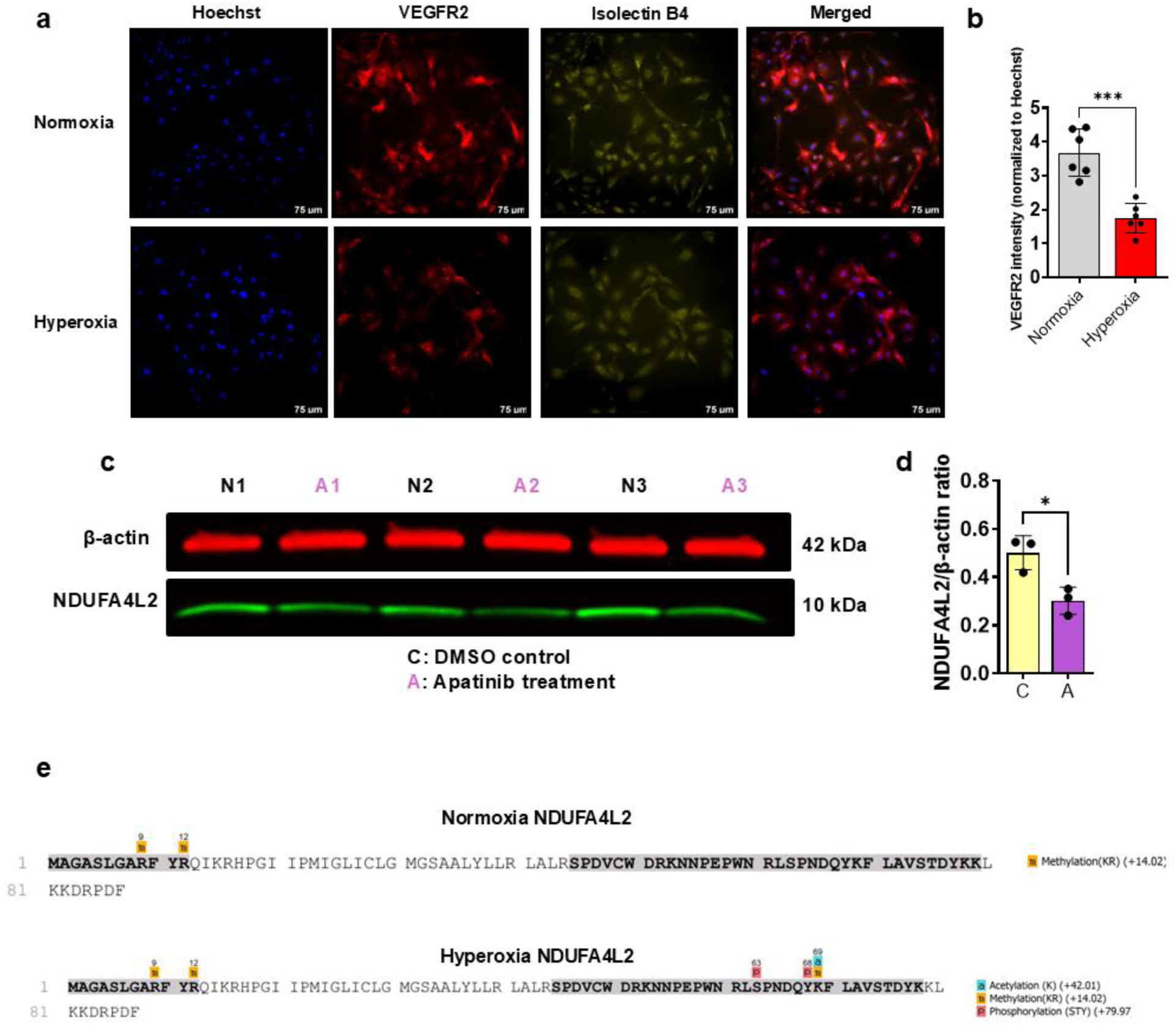
Hyperoxia and Apatinib treatment converge on NDUFA4L2 regulation through a VEGFR2-independent pathway. a) Immunocytochemistry revealed that hyperoxia significantly reduced VEGFR2 expression in primary HRECs. b) quantification of ICC results demonstrated statistically significant differences in VEGFR2 expression. Data are presented as mean ± SD (n = 6 per group). Statistical significance was determined using an unpaired two-tailed Welch’s t-test. c) shows downregulation of NDUFA4L2 by VEGFR2/tyrosine kinase inhibitor, Apatinib, in primary HRECs. d) quantification of western blot resulting from Apatinib-treated or untreated HRECs demonstrated statistically significant downregulation of NDUFA4L2. Data are presented as mean ± SD (n = 3 per group). Statistical significance was determined using an unpaired two-tailed Welch’s t-test. Statistical significance is indicated as: *p- value ≤ 0.05, and ***p-value ≤ 0.001. Legends: C, DMSO control; A, Apatinib treatment. e) PTM analysis showed a unique modification on NDUFA4L2 in hyperoxic conditions.

To directly test whether NDUFA4L2 is regulated by VEGFR2 itself, we first exposed primary HRECs to hyperoxia for 48 hours to reduce basal NDUFA4L2 protein to levels comparable to those at the start of the small-interference (si) RNA experiment, since primary HRECs show high basal NDUFA4L2 expression in normoxia that could mask any further regulation. Cells were then transfected with siRNA against VEGFR2 (siVEGFR2) or non-targeting control (siControl) and treated with or without human VEGF-165 (0 or 50 ng/mL), one of the major isoforms of VEGFA, and then VEGFR2 and NDUFA4L2 protein levels were measured by western blotting. siRNA resulted in roughly 96 percent knockdown (KD) (Supplemental Figure S3a and b). However, siVEGFR2 did not significantly change NDUFA4L2 protein levels compared to siControl (non-targeting controls siRNA), regardless of human VEGF-165 treatment (Supplemental Figure S3a and c). This indicates that NDUFA4L2 protein levels are unaffected by VEGFR2 loss, whether or not cells are concurrently stimulated with VEGF-165, further supporting that VEGFR2 does not directly regulate NDUFA4L2 protein abundance. To further test whether activation, rather than loss, of VEGFR2 regulates NDUFA4L2, we treated serum-starved HRECs with increasing concentrations of human VEGF-165 (20, 50, and 100 ng/mL). Consistent with the siVEGFR2 result, human VEGF-165 (the most abundant VEGFA isoform) treatment did not change NDUFA4L2 protein levels at any concentration tested (Supplemental Figure S3d and e). Together, these results indicate that NDUFA4L2 protein levels are not directly regulated by VEGFR2 signaling, despite the downregulation observed with Apatinib. This implies that there is a previously unknown receptor target of Apatinib that also regulates signaling downstream of VEGFR2, or VEGFR2 KD upregulates other VEGF receptors that compensate for its loss.

We also re-analyzed the published Geo dataset GSE221861 by Rameshekar et al.[48] and found that *NDUFA4L2* transcript is induced in VEGF-treated HRECs, whereas *NDUFA4* shows no change (Supplemental Figure S3f). This suggests that VEGF can regulate NDUFA4L2 at the transcriptional level; however, this transcriptional induction was not reflected in NDUFA4L2 protein abundance in our siVEGFR2 and VEGFA dose-response experiments. This discrepancy implies that VEGF-driven transcription of NDUFA4L2 is uncoupled from its protein levels, and that NDUFA4L2 protein abundance is likely governed by additional, post-transcriptional or post-translational, regulatory mechanisms rather than by VEGFR2 signaling directly.

This also implies that despite the very strong sequence similarities between NDUFA4L2 and NDUFA4, their roles are not redundant, and the regions unique to NDUFA4L2 are potential sites of this additional layer of post-translational regulation. We took the protein sequences from Uniprot and aligned them in Clustal Omega (Supplemental Figure S4a). Interestingly, we found an N-terminal sequence that was unique to NDUFA4L2 (Supplemental Figure S4a). This sequence is known to contain a serine that is phosphorylated in some conditions (source: UniProt). Moreover, the C-terminal sequence also showed some dissimilarities between NDUFA4L2 and NDUFA4, with potential regions of regulation (source: AlphaFold 3; data not shown). Given that NDUFA4L2 protein levels are downregulated by Apatinib but unaffected by direct VEGFR2 KD or VEGFA stimulation, these unique N and C-terminal sequences represent a candidate mechanism through which NDUFA4L2 protein stability could be regulated independently of VEGFR2. To find out if these sites are differentially post-translationally modified (PTM), we performed mass spectrometry analysis of NDUFA4L2 immunoprecipitated from normoxic and hyperoxic cells. Hyperoxic NDUFA4L2 was found to be uniquely modified on S63, Y68, and K69 (Figure 4e). We could not cover hydrophobic regions of the protein in our PTM analysis due to inherent methodological difficulties in measuring hydrophobic regions in small proteins. Since VEGFR2 siRNA inhibition didn’t show a direct regulation of NDUFA4L2, we believe these PTM sites may also be downstream targets of Apatinib or a previously unknown Apatinib-sensitive kinase.

### NDUFA4L2 rescues hyperoxia-mediated migration defects in endothelial cells by rescuing isocitrate dehydrogenase (IDH) flux in the tricarboxylic acid (TCA) cycle

NDUFA4L2 is known to reverse proliferation defects in many cancer models [34, 38–40]. Hyperoxia downregulates both the proliferation and migration of endothelial cells in culture (Figure 5a-c). We first evaluated the effect of NDUFA4L2 on the proliferation of cultured primary HRECs maintained under sub-confluent, actively proliferating conditions. Unlike cancer cells, lentiviral-based NDUFA4L2 overexpression in primary HRECs could not reverse proliferation defects induced by hyperoxia (Figure 5a). We next looked at the effect of NDUFA4L2 overexpression on migration by performing a wound-healing assay on cultured primary HREC stable cell lines at confluency. Interestingly, NDUFA4L2 overexpression successfully rescued hyperoxia-induced migration defects in primary HRECs (Figure 5b and 5c). Since NDUFA4L2 has been shown to inhibit complex I in oxidative phosphorylation in cancer models and activate aerobic glycolysis, which is reasoned to reverse proliferative defects, we tested the effect of NDUFA4L2 overexpression on aerobic glycolysis in our model system. Intracellular metabolites from NDUFA4L2 and mCherry overexpressing cell lines were collected and measured on a mass spectrometer. Intracellular metabolites from NDUFA4L2 overexpressing cells, in sub-confluent cultures used for proliferation assay, showed higher levels of lactate/pyruvate ratio, demonstrating upregulation of aerobic glycolysis in the NDUFA4L2 overexpressing cell line (Figure 5d). It is well established that mitochondria play a critical role in endothelial cell proliferation and migration, functioning as important biosynthetic and signaling hubs that regulate endothelial cell metabolism and adaptation to the local environment, thereby affecting endothelial cell migration, proliferation, and the broader angiogenic process [49]. To further investigate the role of NDUFA4L2 in mitochondrial function, intracellular metabolites from confluent primary HRECs were measured by mass spectrometry. Our untargeted liquid chromatography mass spectrometry (LC-MS) metabolite profiling with an in-house retention time library match showed differences in many TCA cycle metabolites. Aspartate, citrate, and aconitate were found to be increased, whereas α-KG was found to be decreased in hyperoxic cells, indicating a blockage between aconitate and α-KG (Figure 6a and b). Overexpressing NDUFA4L2 rescued this metabolite phenotype and reversed this blockage in confluent primary HRECs exposed to hyperoxia (Figure 6b). We confirmed our findings in an independent gas chromatography mass spectrometry (GC-MS) experiment, which included remaking overexpression lines, exposing them to hyperoxia, and subsequent GC-MS analysis (Figure 6c). These findings suggest that hyperoxia impairs IDH flux—the conversion of isocitrate to αKG—leading to accumulation of citrate and aconitate alongside reduced αKG levels. Interestingly, when we compared these metabolites in sub-confluent primary HRECs to confluent, we did not observe these changes in TCA cycle metabolites (Supplemental Figure S6a). This indicates that the restorative effect of NDUFA4L2 on IDH flux is restricted to migrating cells, rather than proliferating cells.

**Figure 5.**
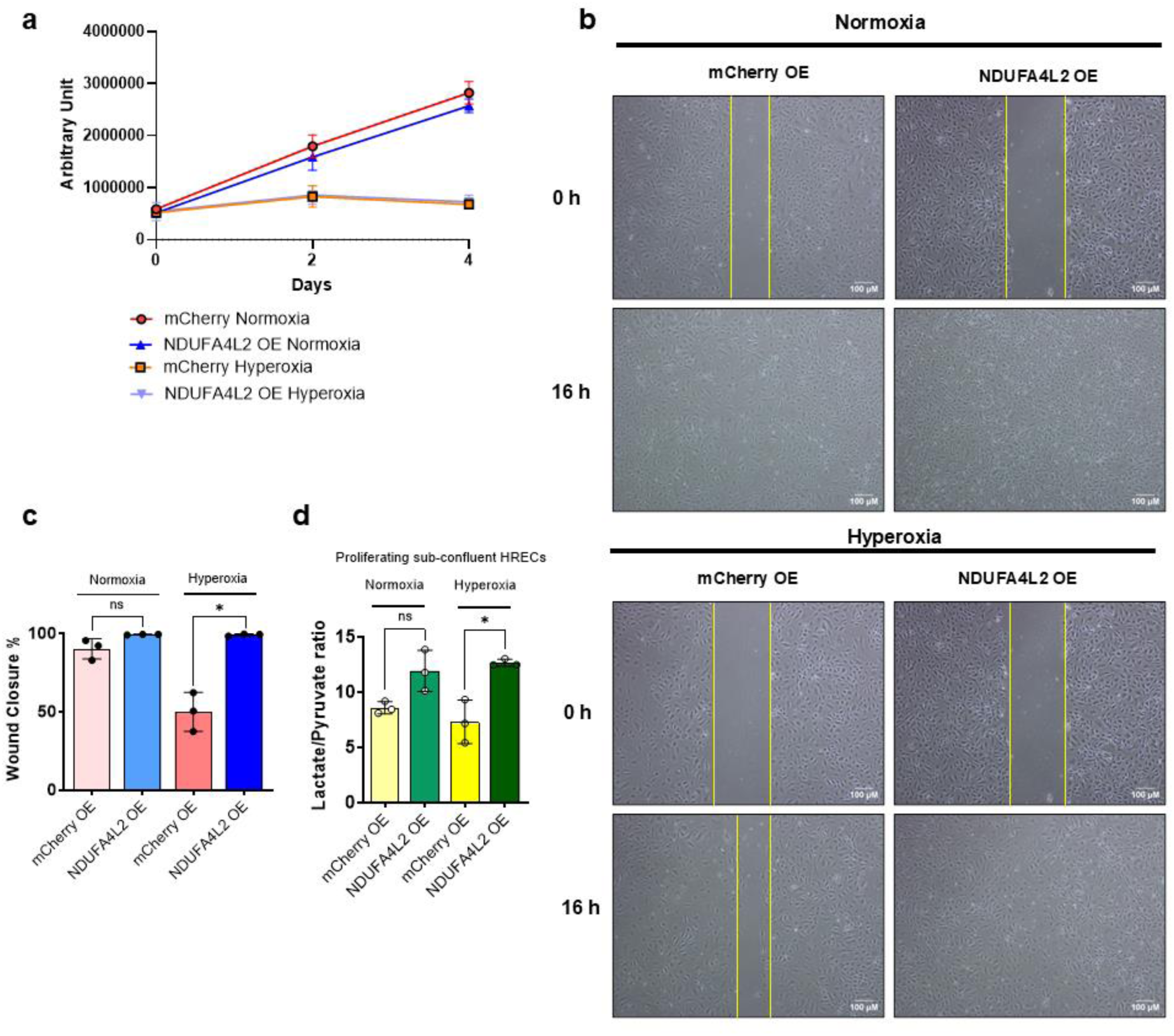
NDUFA4L2 rescues migration but not proliferation phenotype induced by hyperoxia in primary HRECs. a) Unlike cancer cells, in primary HRECs, NDUFA4L2 overexpression didn’t improve or rescue hyperoxia-induced proliferation defect. b) and c) NDUFA4L2 overexpression successfully rescued hyperoxia-mediated inhibition of HREC migration. d) NDUFA4L2 overexpression in sub-confluent HRECs showed increased levels of lactate/pyruvate ratio, indicating upregulation of glycolysis. Data are presented as mean ± SD (n = 3 per group). Statistical significance was determined using an unpaired two-tailed Welch’s t-test. Statistical significance is indicated as: *p-value ≤ 0.05. NS indicates no significant difference.

**Figure 6.**
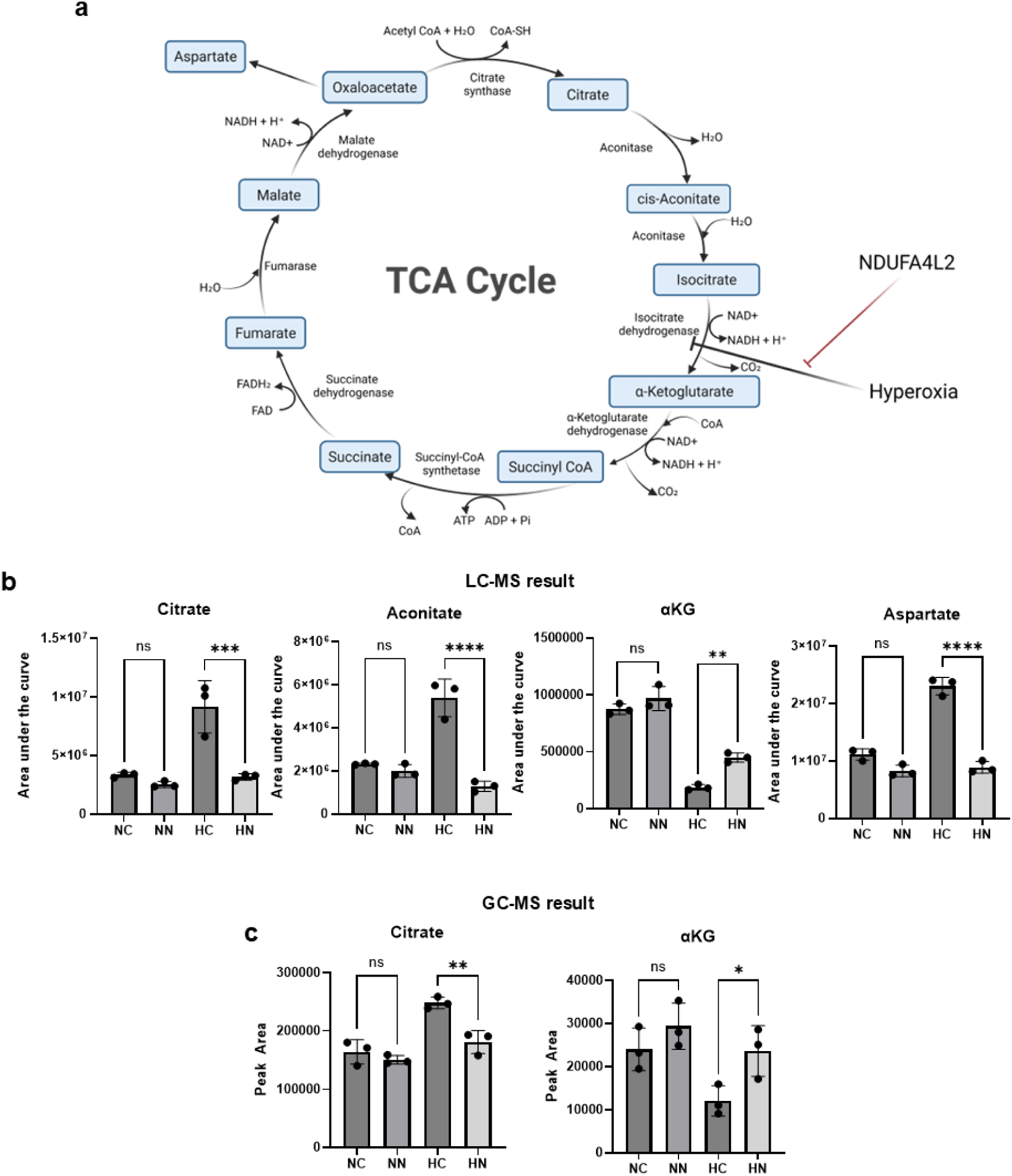
NDUFA4L2 rescues IDH flux impaired by hyperoxia. a) Depicted is a schematic of the tricarboxylic acid (TCA) cycle that shows the potential mechanism of how hyperoxia and NDUFA4L2 impact IDH activity. b) Metabolites upstream of alpha-ketoglutarate (αKG) were found to accumulate in confluent hyperoxia HRECs, whereas αKG was downregulated, implying a blockade in IDH flux. A metabolic phenotypic rescue experiment with NDUFA4L2 overexpression resulted in the recovery of IDH flux. c) An independent GC-MS experiment confirmed the metabolic phenotypic rescue in NDUFA4L2 overexpressing cells. Data are presented as mean ± SD (n = 3 per group). Comparisons among four groups were analyzed using one-way analysis of variance (ANOVA) followed by Tukey’s multiple comparisons post hoc test. Statistical significance is indicated as: *p-value ≤ 0.05, **p-value ≤ 0.01, ***p-value ≤ 0.001 and ****p-value ≤ 0.0001. NS indicates no significant difference. Legends: NC, normoxia mCherry OE; NN, normoxia NDUFA4L2 OE; HC, hyperoxia mCherry OE; HN, hyperoxia NDUFA4L2 OE.

## Materials and methods

### Animal experiments and Oxygen-Induced Retinopathy (OIR) Mouse Model

Animal experiments were performed as per Tufts University and Massachusetts General Hospital Institutional Animal Care and Use Committee-approved protocols. We followed OIR protocol as previously published by Smith et al. [50]. For OIR experiments, mice were kept in room air from postnatal day (P) 0 to P7. On P7, the hyperoxic treatment group was moved to 75% oxygen, whereas the control group was kept in room air until P12, leading to phase I or vaso-obliteration of OIR. Euthanasia was achieved by treating mice with Isoflurane followed by cervical dislocation. All the mice were anesthetized with isoflurane and euthanized by cervical dislocation before the removal of tissues. At P12, retinas were harvested on ice. For protein extraction, retinas were rinsed with phosphate-buffered saline (PBS) and homogenized in 250 µL RIPA lysis buffer (Boston BioProducts, Milford, MA, USA) supplemented with protease and phosphatase inhibitors (Roche, Basel, Switzerland). Tissue lysates were briefly sonicated and centrifuged at 20,000 × g for 15 min at 4 °C. Supernatants were collected and stored at −80 °C until analysis.

### Cell Culture

Primary human retinal endothelial cells (HRECs) were obtained from Cell Systems (Kirkland, WA, USA) and used between passages 4 and 5. Cells were maintained in attachment factor-coated plates (Cell Systems; catalog no. 4Z0-201) with human endothelial cell growth medium with endothelial cell medium supplement kit (Cell Biologics, Chicago, IL, USA; catalog no. H1168)

### Hypoxia-inducible Factor 1 alpha (HIF1α) Stabilization by Roxadustat

Primary HRECs were seeded in attachment factor–coated 6-well plates (Cell Systems) and maintained in human endothelial growth medium (Cell Biologics). For experiments, cells were plated at a density of 0.1 × 10⁶ cells per well and allowed to adhere for 24 h. Following attachment, cells were exposed to either normoxic (21% O₂) or hyperoxic (75% O₂) conditions for a total of 96 h. Roxadustat (FG-4592) was purchased from Selleck Chemicals (Selleck Chemicals, Houston, TX, USA; catalog no. S1007), and 50 mg/mL stock was prepared in dimethyl sulfoxide (DMSO). The stock was further diluted with PBS to make a 1 mg/mL working stock. After 72 h of oxygen exposure, cells were treated with either vehicle control (diluted DMSO in PBS; 20 µL in 2 mL media) or 10 µg/mL Roxadustat for an additional 24 h. At the end of the treatment period, cells were harvested for protein and RNA analyses. Detailed procedures for protein extraction and RNA isolation are described in the sections below (“Protein Extraction from Cultured Cells” and “RNA Isolation and Quantitative RT-PCR”).

### HIF1α Stabilization by MG-132

Primary HRECs were seeded in attachment factor–coated 6-well plates (Cell Systems) at a density of 0.1 × 10⁶ cells per well and allowed to adhere for 24 h. Following attachment, cells were maintained under either normoxic (21% O₂) or hyperoxic (75% O₂) conditions for a total of 96 h. After 90 h of oxygen exposure, cells were treated with either vehicle control (DMSO; 2 µL in 2 mL media) or 10 µM MG-132 (Sigma-Aldrich, St. Louis, MO, USA; catalog no. M7449- 200UL) for the final 6 h. Cells were harvested immediately after treatment for protein and RNA analyses. Detailed procedures for protein extraction and RNA isolation are described in the sections below (“Protein Extraction from Cultured Cells” and “RNA Isolation and Quantitative RT-PCR”).

### Apatinib treatment in HRECs

Primary HRECs were seeded in attachment factor (Cell System)–coated 6-well plates and maintained in human endothelial growth medium (Cell Biologics). For experiments, cells were plated at a density of 0.1 × 10⁶ cells per well and allowed to adhere for 24 h. Following attachment, cells were treated with 20 µM Apatinib (a VEGFR2 tyrosine kinase inhibitor, Cayman Chemical, Ann Arbor, MI, USA: catalog no.21268) for 72 h to assess whether VEGFR2 inhibition downregulates NDUFA4L2 protein expression. At the end of the treatment period, cells were harvested for protein extraction and analyzed by western blotting.

### Human Vascular Endothelial Growth Factor (VEGF)-165 dose-dependent treatment of HRECs

Primary HRECs were seeded at a density of 0.15 × 10⁶ cells per well in 6-well plates and incubated for 24 h at 37°C in 5% CO₂ in complete human endothelial cell growth medium (Cell Biologics) to allow attachment. Cells were then serum-starved for 8 h in high-glucose DMEM (Dulbecco’s Modified Eagle Medium) supplemented with 1× sodium pyruvate and 1× GlutaMAX (Thermo Fisher Scientific) to synchronize cells prior to human VEGF-165 stimulation.

A human VEGF-165 working stock (10 µg/mL) was freshly prepared by reconstituting lyophilized recombinant human VEGF-165 (Thermofisher Scientific) in 0.3% bovine serum albumin (BSA) in PBS, and used immediately for treatment. Serum-starved HRECs were treated with VEGFA at final concentrations of 20, 50, or 100 ng/mL, respectively, in 2 mL of complete human endothelial cell growth medium (Cell Biologics). Cells were incubated with VEGF-165 at 37°C in 5% CO₂ for 4 h. At the end of each treatment period, cells were washed, and whole-cell protein lysates were collected for downstream analysis of NDUFA4L2 protein levels by western blotting.

### Small-interference (si) RNA-mediated knockdown of VEGF Receptor2 (VEGFR2)

Primary HRECs were seeded in attachment factor–coated 6-well plates (Cell Systems) and maintained in human endothelial growth medium (Cell Biologics). For experiments, cells were plated at a density of 0.1 × 10⁶ cells per well. Following attachment, cells were transferred to a hyperoxic chamber (75% O₂, 5% CO₂) for 48 h. HRECs were transfected with small interfering RNA (siRNA) targeting VEGFR2 (siVEGFR2) or a non-targeting control siRNA (siCtrl) using Lipofectamine 3000 (Thermo Fisher Scientific), according to the manufacturer’s protocol with minor modifications. Lyophilized siRNA (Horizon Discovery, Lafayette, CO, USA) was resuspended in nuclease-free UltraPure water to a stock concentration of 50 µM and stored on ice until use. Prior to transfection, the growth medium was aspirated and replaced with 1 mL of Opti-MEM Reduced Serum Medium (Thermo Fisher Scientific) per well of a 6-well plate. Transfection complexes were prepared by combining siRNA and Lipofectamine 3000 in separate tubes. In the first tube, siRNA was diluted in Opti-MEM. In the second tube, Lipofectamine 3000 was diluted in Opti-MEM and incubated for 5 min at room temperature. The diluted siRNA was then combined with the diluted Lipofectamine 3000, followed by the addition of Opti-MEM, and the mixture was incubated for 15 min at room temperature to allow transfection complex formation. The resulting transfection mixture was added, and cells were incubated at 37°C in 5% CO₂ for 6 h. After 6 h of incubation, the transfection medium was replaced with endothelial growth medium (Cell Biologics) supplemented with or without human VEGF-165 (Thermofisher Scientific, prepared as described above) at a final concentration of 50 ng/mL and incubated overnight at 37 °C in 5% CO₂. At the end of the treatment period, cells were harvested for downstream analyses.

### Construction of Plasmids for mCherry or human NDUFA4L2 Overexpression

Two lentiviral expression constructs were generated using the pLJM1-empty backbone (Addgene, Watertown, MA, USA; plasmid #91980) [51], which was digested with NheI-HF (New England Biolabs, Ipswich, MA, USA) to facilitate insert cloning, according to the manufacturer’s instructions. The coding sequences of mCherry or human NDUFA4L2 were PCR-amplified from pcDNA3.1 plasmids (GenScript, Piscataway, NJ, USA) containing the respective inserts, using the primers listed below and the NEBNext® High-Fidelity 2× PCR Master Mix (New England Biolabs). Primers were designed to be compatible with the NheI-HF–digested vector ends. PCR amplification was performed with an initial denaturation at 98 °C for 30 s, followed by 35 cycles of denaturation at 98 °C for 10 s, annealing at 72 °C for 30 s, and extension at 72 °C for 22 s (mCherry) or 8 s (human NDUFA4L2), with a final extension at 72 °C for 2 min. The amplified PCR products were separated on a 1.5% agarose gel, and bands corresponding to the expected size were excised and purified using the QIAquick Gel Extraction Kit (QIAGEN, Germantown, MD, USA) according to the manufacturer’s protocol. Purified PCR products were assembled into the linearized pLJM1- empty vector using a Gibson assembly–based approach with the NEBuilder HiFi DNA Assembly Kit (New England Biolabs), according to the manufacturer’s instructions. The assembled plasmids were transformed into NEB® Stable Competent Escherichia coli (High Efficiency) (New England Biolabs). Following transformation, individual colonies were selected, and plasmid constructs were sequence-verified by Plasmidsaurus (Plasmidsaurus Inc., Louisville, KY, USA).

### Cloning primers

mCherry

Forward: TTA GTG AAC CGT CAG ATC CGA TGG TGA GCA AGG GCG AG

Reverse: CGG ACT TGT ACC CGG TAG CGC TAC TTG TAC AGC TCG TCC ATG

Human NDUFA4L2

Forward: TTA GTG AAC CGT CAG ATC CGA TGG CAG GAG CCA GTC TTG

Reverse: CGG ACT TGT ACC CGG TAG CGT TAG AAG TCT GGC CGG TC

### Lentivirus Production

Lentiviral particles encoding human NDUFA4L2 or mCherry were generated by transient triple transfection of 293-FT cells (Thermo Fisher Scientific, Waltham, MA, USA) using pCMV-VSV-G (Addgene; plasmid #8454)[52], psPAX2 (Addgene; plasmid #12260), and either pLJM1-NDUFA4L2 or pLJM1-mCherry plasmids. Briefly, 300 ng of pCMV-VSV-G, 2,700 ng of psPAX2, and 3,000 ng of each expression construct (pLJM1-NDUFA4L2 or pLJM1-mCherry) were mixed in 500 µL of Opti-MEM™ (Thermo Fisher Scientific). Subsequently, 18 µL of X- tremeGENE™ HP DNA Transfection Reagent (Roche, Basel, Switzerland) was added, and the mixture was incubated for 20 min at room temperature. The transfection complex was then added to 293-FT cells at approximately 70% confluency, cultured in 100 mm dishes. Viral supernatants were collected 48 h post-transfection and filtered through 0.45 µm PES syringe filters (CELLTREAT, Ayer, MA, USA).

### Generation of Stable HREC Lines

HRECs at approximately 70% confluency were transduced with mCherry or NDUFA4L2 lentivirus in the presence of polybrene (Sigma-Aldrich; 1:1000 dilution). After 48 h, viral media were replaced with fresh human endothelial growth medium (Cell Biologics). Transduced cells were selected using puromycin (2 µg/mL) for 48 h. Stable cell lines were replated at 0.1 × 10⁶ cells per well in attachment factor (Cell System)-coated 6-well plates and, after attachment, exposed to either normoxic or hyperoxic conditions for 48 h. Spent media and cell lysates were subsequently collected.

### Protein Extraction from Cultured Cells

Cells were washed with PBS and lysed in 80 µL RIPA buffer (Boston BioProducts) containing protease and phosphatase inhibitors (Roche, Basel, Switzerland). Lysates were collected by scraping, briefly sonicated, and centrifuged at 20,000 × g for 15 min at 4 °C. Cleared supernatants were stored at −80 °C until use.

### SDS-PAGE and Western Blot Analysis

Protein concentrations were determined using the Pierce™ BCA Gold Protein Assay (Thermo Fisher Scientific). Equal amounts of protein (15–20 µg) were mixed with Tris-glycine SDS sample buffer supplemented with 20 mM DTT, heated at 95 °C for 5 min, and briefly centrifuged. Samples were resolved on either 4–20% Novex™ WedgeWell™ Tris-glycine precast gels (Invitrogen, Waltham, MA, USA) or 4–15% Mini-PROTEAN® TGX™ precast gels (Bio-Rad Laboratories, Hercules, CA, USA). Electrophoresis was performed at 200 V for Novex gels and 120 V for TGX gels. Proteins were transferred to 0.45-µm PVDF membranes (MilliporeSigma, Burlington, MA, USA) by wet transfer in Tris-glycine buffer at 30 V overnight in a cold room.

Membranes were air-dried for 1 h, briefly activated with methanol, rinsed with water, and equilibrated in TBS. Blots were blocked with Intercept® TBS Blocking Buffer (LI-COR Biosciences, Lincoln, NE, USA) for 1 h at room temperature and incubated with primary antibodies diluted in blocking buffer containing 0.2% Tween-20 overnight at 4 °C. After washing with TBST (three washes, 10 min each), membranes were incubated with IRDye®- conjugated secondary antibodies diluted in blocking buffer containing 0.2% Tween-20 and 0.01% SDS for 1 h at room temperature in the dark. Blots were washed with TBST and rinsed with TBS. Signals were captured using a ChemiDoc™ MP Imaging System (Bio-Rad Laboratories). Western blot band intensities were quantified using Image Lab software version 6.1 (Bio-Rad Laboratories, Hercules, CA, USA), and protein expression levels were normalized to the internal loading control, β-actin.

### Primary antibodies

NDUFA4L2: rabbit polyclonal antibody, catalog # 16480-1-AP (Proteintech, Rosemont, IL, USA).

NDUFA4L2: rabbit polyclonal antibody, catalog # PA5-106688 (Thermo Fisher Scientific, Waltham, MA, USA).

HIF-1α (D1S7W): rabbit monoclonal antibody, catalog # 36169S (Cell Signaling Technology, Danvers, MA, USA).

Hydroxy-HIF-1α (Pro564) (D43B5): rabbit monoclonal antibody, catalog # 3434S (Cell Signaling Technology, Danvers, MA, USA).

β-actin: (8H10D10) mouse monoclonal antibody, catalog # 3700S (Cell Signaling Technology, Danvers, MA, USA).

VEGFR2 (D5B1): rabbit monoclonal antibody, catalog # 9698T (Cell Signaling Technology, Danvers, MA, USA).

### Secondary antibodies

IRDye® 800CW donkey anti-rabbit IgG (H+L), polyclonal, catalog # 925-32213 (LI-COR Biosciences, Lincoln, NE, USA).

IRDye® 680RD donkey anti-mouse IgG (H+L), polyclonal, catalog # 925-68072 (LI-COR Biosciences, Lincoln, NE, USA).

Secondary antibodies were used at a 1:2000 dilution.

### RNA Isolation and Quantitative Reverse Transcription Polymerase Chain Reaction (qRT-PCR)

Total RNA was isolated using TRI reagent (Sigma-Aldrich) according to the manufacturer’s instructions. cDNA synthesis was performed using the iScript™ gDNA Clear cDNA Synthesis Kit (Bio-Rad). 2 µL of this cDNA was mixed with 5 µL of 2X iTaq™ Universal SYBR® Green Supermix (Bio-rad), 0.5 µL of 10 µM forward primer, 0.5 µL of 10 µM reverse primer, and 2 µL of nuclease-free water. Amplification was performed with an initial incubation at 50 °C for 2 min, followed by an initial denaturation at 95 °C for 10 min. PCR amplification was then carried out for 40 cycles of 95 °C for 15 s and 60 °C for 1 min, with fluorescence acquisition at the end of each extension step. Melt curve analysis was subsequently performed with a denaturation step at 95 °C for 15 s, followed by a gradual increase from 65 °C to 95 °C with 0.5 °C increments and continuous fluorescence acquisition.

### qRT-PCR primers

Human ANGPTL4

Forward: TAC CCT TCT CCA CTT GGG AC

Reverse: AAA CCA CCA GCC TCC AGA GA

Human PDK1

Forward: AGT TCC TGG ACT TCG GAT CA

Reverse: AAC GGA TGG TGT CCT GAG AA

Human NDUFA4L2

Forward: GAT CGG CTT AAT CTG CCT GG

Reverse: CGG GTT GTT CTT TCT GTC CC

### Cell Proliferation Assay

HREC stable cell lines (mCherry overexpression (OE) and human NDUFA4L2 OE) were seeded in attachment factor (Cell Systems)–coated black-walled, clear-bottom 96-well plates at a density of 2,000 cells per well and maintained in human endothelial growth medium (Cell Biologics). Cells were allowed to adhere for 24 h, after which baseline cell number (Day 0) was measured using the CyQUANT™ NF Cell Proliferation Assay (Thermo Fisher Scientific) according to the manufacturer’s instructions. Cells were then exposed to normoxic (21% O₂) or hyperoxic (75% O₂) conditions for 4 days, with media replaced every 2 days. Cell proliferation was assessed on Day 0, Day 2, and Day 4 using the CyQUANT™ NF Cell Proliferation Assay.

### Wound-Healing Assay

HREC stable cell lines (mCherry OE and human NDUFA4L2 OE) were plated in attachment factor (Cell Systems)-coated 6-well plates with full confluency and then exposed to normoxic or hyperoxic conditions for 24 h. A linear scratch was created using a sterile 1-mL pipette tip, followed by 3 times washing with PBS to remove detached cells. Cells were incubated in human endothelial growth medium (Cell Biologics) for an additional 24 h, exposed to normoxic or hyperoxic conditions. Images were captured immediately after scratching and at 16 h. Wound closure % was quantified using the Wound Healing Tool plugin in ImageJ software (NIH, USA).

### Metabolite Extraction from Cell Culture Media

HREC stable cell lines (mCherry OE and human NDUFA4L2 OE) were plated in attachment factor (Cell Systems)-coated 6-well plates with 70% or full confluency and then exposed to normoxic or hyperoxic conditions for 48 h. Spent media collected from stable cell lines were stored at −80 °C. For metabolite extraction, 40 µL of media was mixed with 160 µL 80% methanol with a final concentration of 0.5 mM of tricarballylic acid (TCLA) (Sigma-Aldrich), as an internal standard. Samples were incubated at −80 °C for 1 h. Samples were vortexed and centrifuged at 15,000 × g for 5 min at 4 °C. 50 µL of supernatant was transferred to new tubes and dried using a SpeedVac concentrator for derivatization.

### Intracellular Metabolite Extraction

For intracellular metabolite extraction, cells were washed with 1 mL of normal saline. The saline was then aspirated, and 1 mL of ice-cold 80% methanol with a final concentration of 0.02 mM TCLA (Sigma-Aldrich) was added to each well. Cells were scraped on ice using a cell scraper, collected into 1.5 mL tubes, and centrifuged at 15,000 × g for 5 min at 4 °C. Subsequently, 500 µL of the supernatant was transferred to new tubes and dried using the SpeedVac concentrator for derivatization.

### Sample Derivatization and Gas Chromatography–Mass Spectrometry (GC-MS) Based Metabolomics

Dried samples were resuspended in 25 µL methoxyamine hydrochloride (40 mg/mL in pyridine) and incubated at 45 °C for 30 min with shaking. Following cooling to room temperature, 25 µL MSTFA+1%TMCS (Macherey-Nagel, Düren, Germany) was added, and samples were incubated at 45 °C for an additional 30 min. Samples were centrifuged at 15,000 × g for 2 min, and 60 µL of supernatant was transferred to GC-MS vials with inserts for analysis.

Derivatized Samples were run on an Agilent 8890 gas chromatograph interfaced with a 5977B EI/CI mass selective detector, operating in electron impact mode with an extractor- type ion source. One μL of sample was introduced via split injection at a 3:1 split ratio, directing a portion of the sample into the mass spectrometer. Mass spectral data were acquired in full-scan mode over a range of 70–700 m/z. The inlet heater was maintained at 250 °C, with a septum purge flow of 3 mL/min. Chromatographic separation was achieved using a DB-5ms capillary column (30 m × 250 μm × 0.25 μm) coupled to the MSD, operated under constant helium flow at 1 mL/min. The oven program began at 60 °C, held for 1 min, then increased at a rate of 10 °C/min up to 335 °C, where it was held for an additional 5 min to ensure complete analyte separation.

### Metabolite extraction and Liquid Chromatography–Mass Spectrometry (LC-MS) Based Metabolomics

HREC stable cell lines (mCherry OE and human NDUFA4L2 OE) were plated in attachment factor (Cell Systems)-coated 6-well plates with full confluency and then exposed to normoxic or hyperoxic conditions for 24 h. Labeling medium was prepared fresh immediately prior to use. Briefly, a 20 mM L-glutamine stock solution was prepared in glucose-free DMEM (no additives). A separate 7.5 mM uniformly labeled ¹³C₆-glucose stock solution was prepared in glucose-free DMEM. The two stock solutions were then combined by mixing 9.5 mL of the ¹³C₆-glucose stock with 500 µL of the L-glutamine stock. The combined solution was further diluted by adding 9 mL of this mixture to 18 mL of standard HREC growth medium, yielding a final labeling medium with a ¹³C₆-glucose concentration of 2.5 mM and an L-glutamine concentration of 0.33 mM. Cells in both normoxic and hyperoxic chambers were then incubated with this labeling medium for an extra 24 hours prior to metabolite extraction.

Untargeted metabolomics was performed on a Vanquish Horizon LC system coupled to an Exploris 480 Orbitrap mass spectrometer (Thermo Fisher Scientific). Chromatographic separation was performed by hydrophilic interaction liquid chromatography (HILIC) on an ACQUITY UPLC BEH amide column (Waters Corporation, Milford, MA, USA; P/N 186004802; 2.1 × 150 mm, 1.7 μm, 130 Å). Mobile phase A consisted of 95:5 H₂O:acetonitrile with 10 mM ammonium acetate and 10 mM ammonium hydroxide, and mobile phase B consisted of 20:80 H₂O:acetonitrile with 10 mM ammonium acetate and 10 mM ammonium hydroxide. The gradient was as follows: 100% B from 0.0–3.0 min, 90% B from 3.2–6.2 min, 80% B from 6.5–10.5 min, 70% B from 10.7–13.5 min, 45% B at 13.7–16 min, and re-equilibration to 100% B from 16.5–22 min. Injection volume was 5 μL. Mass spectrometric detection was performed in full scan mode with polarity switching, over a scan range of 70–1000 m/z, at an Orbitrap resolution of 120,000.

### Retinal flatmount immunohistochemistry

Immunostaining of retinal flatmounts was performed as described briefly [53]. In brief, eyes collected from P12 C57BL6J pups were fixed in phosphate-buffered 4% paraformaldehyde (PFA) solution (Electron Microscopy Sciences, Morgantown, PA) for 10-20 minutes at room temperature. Following fixation, the eyes were dissected in 1X-PBS and the retinal flatmounts post-fixed in ice-cold 100% methanol and stored at 4 °C in 100% methanol until staining. Prior to staining the retinal flatmounts were washed in 1X-PBS, blocked overnight in blocking buffer containing 10% fetal bovine serum and 0.05% triton-X100 in 1X PBS. The blocked retinal flatmounts were incubated with NDUFA4L2 antibody (1:400, cat# PA5- 106688, Thermo Fisher Scientific) for 36-48 hr at 4 °C. The retinal flatmounts were washed extensively in 1X-PBS and incubated overnight at 4 degree 4 C with donkey-anti rabbit, Alexa Fluor™ 555 (1:500, cat# A-31572, Thermo Fisher Scientifi), Isolectin GS-IB4, Alexa Fluor™, 647 conjugate (1:250, cat# I32450, Thermo Fischer Scientific, Waltham, MA), with or without alpha smooth muscle actin (1A4) Alexa Fluor™ 488 (5µg/ml, cat# 53-9760-82, Thermo Fisher Scientific). After thorough washes in 1X-PBS, retinal flatmounts were mounted onto the slides using Prolong Gold anti-fade reagent (cat# P36934, Thermo Fisher Scientific). The slides were then imaged using a Leica Thunder fluorescence microscope and analyzed.

### Immunocytochemistry for VEGFR2 in HRECs

Primary HRECs were seeded at a density of 0.5 × 10⁵ cells per well into chamber slides pre-coated with attachment factor (Cell Systems), with three wells per group (normoxia and hyperoxia). Following overnight incubation, the medium was replaced with fresh growth medium, and chambers were placed under either normoxic (21% O₂) or hyperoxic (75% O₂) conditions for 48 h.

Cells were washed twice with warm PBS and fixed with 1 mL of 4% PFA in PBS (Boston BioProducts) for 15 min at room temperature. Following fixation, cells were washed twice with PBS and permeabilized with 1 mL of 1% Triton X-100 (Sigma-Aldrich) for 30 min at room temperature. Cells were then blocked overnight at 4°C in 0.3% Triton X-100 containing 3% bovine serum albumin (BSA, w/v).

The following day, cells were washed twice with PBS and incubated overnight at 4°C with primary antibody against VEGFR2 (Cell Signaling Technology, catalog# 9698S) diluted 1:300 in 250 µL of 0.3% Triton X-100 containing 3% BSA (w/v). Cells were subsequently washed three times for 5 min each with PBS at room temperature and incubated for 2 h at room temperature with a secondary antibody cocktail consisting of Alexa Fluor 488-conjugated Goat anti-rabbit IgG (1:1000) (Thermofisher Scientific, catalog# A11008), Isolectin GS- IB4 From *Griffonia simplicifolia*, Alexa Fluor™ 568 Conjugate (1:500) (Thermofisher, catalog# I21412), and Hoechst 33258 nuclear stain (1 µg/mL, diluted from a 10 mg/mL stock) (Thermofisher Scientific, catalog# H3569). Cells were then washed three times for 5 min each with PBS at room temperature, mounted with VECTASHIELD PLUS antifade mounting medium (Vector Laboratories, Newark, CA, USA; Cat# H-1900), and imaged with the Leica Thunder fluorescence microscope. A total of 6 images were taken from each group and the expression was quantified using ImageJ software (NIH, USA).

### Protein extraction, tryptic digestion, Tandem Mass Tag (TMT) labelling and Liquid Chromatography-Tandem Mass Spectrometry (LC-MS/MS) analysis

Retina tissue was lysed in 1X SDS lysis buffer (5 % (w/v) SDS, 100mM triethylammonium bicarbonate (TEAB), pH 8.5, 40mM CAA, 10mM tris(2-carboxyethyl)phosphine (TCEP)) using a pestle for mechanical disruption. Lysates were heated at 90 °C and extracted via the SP3 method as in Hughes et al. in 2019 [54]. Proteins were bound to SP3 beads by adding 100 % (v/v) ethanol, followed by three washing steps with 80 % (v/v) ethanol. Proteins were digested with trypsin/LysC mix (1:100) in 100mM TEAB overnight in a shaking incubator at 37 °C at 115 RPM. The following day, an additional dose of trypsin/Lys-C mix was added (1:100) in 100mM TEAB and the digestion continued for 4 hours at 37 °C. Peptide digests were dried in a speed- vac concentrator and resuspended in 100mM TEAB. Peptides were labeled with TMT resuspended in anhydrous acetonitrile. TMT-labeled samples were quenched with 5% hydroxylamine in 100mM TEAB. Labeled peptides were pooled together and purified using EasyPep columns (Thermo Scientific) and fractionated using a Strong Anion Exchange column (Thermo Scientific). Increasing concentrations of ammonium acetate (0, 20, 50, 100, 200, 500 mM) were used for elution. Low salt fractions (0, 20, 50 mM ammonium acetate) and high salt fractions (100, 200, 500 mM ammonium acetate) were pooled respectively and lyophilized. The High pH Reversed-Phase Peptide Fractionation Kit (Thermo Scientific) was used following a 12-step gradient of increasing acetonitrile concentrations: 5, 7.5, 10, 12.5, 15, 17.5, 20, 22.5, 25, 27.5, 30, 60%. The following fractions were then pooled together and lyophilized: 1+7, 2+8, 3+9, 4+10, 5+11, 6+12. Peptide fractions were resuspended in 0.2% formic acid in MS-grade water for LC-MS analysis.

Mass spectrometry was performed using an Orbitrap Exploris 480 mass spectrometer connected to an Easy-nLC 1200 chromatography system using an EasySpray ES900 column (75 µm x 15 cm, 100 Ȧ), all from Thermo Fisher Scientific. Peptides were separated at 300 nl/min on a gradient of 1–25% B over 90 min, 25–40% B over 30 min, 40–95% B over 10 min, 95% B for 10 min, 95-2% B over 2 min, 2% B for 2 min, 2-98% B over 2 min, 98% B for 2 min, 98-2% B over 2 min, 2% B for 2 min, using 0.1% FA in water for A and 0.1% FA in 80% acetonitrile for B. The Orbitrap was operated in positive ion mode with a positive ion voltage of 1800 V, an ion transfer tube temperature of 270 °C, and a 4.6 L/min carrier gas flow. Spectra were acquired at a resolution of 120,000 (MS1) and 30,000 (MS2), with a scan range of 350–1200 m/z, standard AGC target, isolation windows of m/z 0.7, intensity threshold of 5.0e3, 2–5 charge state, dynamic exclusion of 30 seconds, and 36% HCD collision energy. The Fragpipe 21.1 software package was used to analyze proteome data following the methodology outlined by Schulte et al. in 2019 [55]. For protein quantification, raw protein peak areas were calculated with at least two highly abundant peptide signals.

### NDUFA4L2 Immunoprecipitation (IP)

NDUFA4L2 overexpressing stable HRECs were seeded at 0.5 × 10⁶ cells per 100 mm dish and maintained under either normoxic (21% O₂) or hyperoxic (75% O₂) conditions for 48 h. Cells were washed with ice-cold PBS and lysed in 300 µL of IP lysis buffer (20 mM Tris, 150 mM NaCl, 0.5 % IGEPAL CA-630, pH 8) supplemented with protease and phosphatase inhibitors (Cell Signaling Technology). Lysates were incubated on ice for 15 min, sonicated, and clarified by centrifugation at 14,000 × g for 10 min at 4 °C. 200 µL of clarified lysate was incubated with 5 µL of anti-NDUFA4L2 antibody (Thermo Fisher Scientific) overnight at 4 °C with gentle rotation. Immunocomplexes were captured using pre-washed magnetic Protein A beads (Cell Signaling Technology) for 1 h at 400 rpm, followed by five washes with 500 µL of lysis buffer. Beads were then resuspended in 40 µL of IP lysis buffer containing protease and phosphatase inhibitors (Cell Signaling Technology).

For elution, immunoprecipitated proteins were mixed with 2× Novex™ Tris-Glycine SDS Sample Buffer (Thermo Fisher Scientific) and 10× NuPAGE™ Sample Reducing Agent (Thermo Fisher Scientific), and heated at 90 °C for 5 min. Magnetic beads were removed using a magnetic separation rack, and the resulting eluates were stored at −80 °C until post-translational modification (PTM) analysis.

### Post-Translational Modification (PTM) analysis

Samples underwent solubilization, reduction, and alkylation for 15 min at 70 °C using 5% (w/v) SDS and 100 mM TEAB supplemented with 40 mM CAA and 10 mM TCEP. Protein extraction was performed using the SP3 method, following the protocol of Hughes et al. (2019). Overnight digestion was carried out with trypsin/Lys-C mix (1:100) in 100 mM TEAB at 37 °C in a shaking incubator set to 115 RPM. On the following day, a second aliquot of trypsin/Lys-C mix (1:100) in 100 mM TEAB was added, and digestion proceeded for an additional 4 hours at 37 °C. Resulting peptide digests were purified using stage tips according to the method of Rappsilber et al. (2007)[56]. Peptides were subsequently dried using a speed-vac concentrator at 55 °C, then resuspended in 0.2% (v/v) formic acid in MS-grade water prior to LC-MS analysis. LC-MS/MS measurements were conducted using a Vanquish Neo nanoLC system interfaced with an Orbitrap Eclipse mass spectrometer, equipped with a FAIMS Pro Interface and an Easy Spray ESI source (all Thermo Fisher Scientific). Chromatographic separation was achieved using an Aurora Ultimate XT25 column (75 µm × 25 cm, 120 Å; IonOpticks, Fitzroy, VIC, Australia). The mobile phase comprised 0.1% (v/v) formic acid in water (solution A) and 0.1% (v/v) formic acid in 80% (v/v) acetonitrile (solution B), delivered at a flow rate of 400 nL/min with the column maintained at 40 °C. The elution gradient consisted of 3-25% B over 90 min, 25-40% B over 30 min, 40- 95% B over 10 min, and a final hold at 95% B for 6 min.

The mass spectrometer was operated in positive ion mode with the ion source temperature set to 305 °C; ionized peptides passed through the FAIMS Pro unit at applied voltages of -45, -55, and -65 V. MS1 spectra were acquired at a resolution of 120,000 across a mass range of 375–1500 m/z, using a custom AGC target of 100 and automatic injection time. MS2 acquisition was performed in DDA mode with dynamic exclusion enabled and a resolution of 15,000. Precursors with charge states of 2-7 were selected for MS2 analysis, applying a 60-second dynamic exclusion window. MS2 parameters included dynamic maximum injection time, standard AGC target, a 1.6 m/z isolation window, 30% normalized collision energy, and an intensity threshold of 2.5E4.

Data analysis was performed with PEAKS Studio (version 10.6). Raw files were processed using a retention time tolerance of 10 ppm and a precursor mass window of 10 ppm, considering precursor charge states from +2 to +8. Preprocessing steps included automatic centroiding, deisotoping, and deconvolution of MS spectra. Protein identification employed the NDUFA4L2 sequence as a reference. Search parameters specified a parent ion mass tolerance of 15 ppm and a fragment ion mass tolerance of 0.05 Da, with trypsin designated as the protease allowing specific cleavage (D/P) and up to three missed cleavage sites. Methionine oxidation, phosphorylation, acetylation, and ubiquitination were set as variable modifications, cysteine carbamidomethylation was set as a fixed modification, with a maximum of three variable modifications permitted per peptide. FDR was estimated and controlled using a decoy-fusion strategy, with peptide and protein identifications filtered to a significance threshold of -10logP ≥ 20. Quantified data were exported as .csv files and analyzed further according to the approach described by Schulte et al. [57].

### Statistical analysis

Normalized TMT-labeled proteomics data analysis was performed on metaboanalyst.ca.

Statistical analyses were performed using GraphPad Prism version 10 (GraphPad Software, San Diego, CA, USA). Data are presented as the mean ± standard deviation (SD). Comparisons between two independent groups were performed using Welch’s unpaired *t*- test. Comparisons among three or more groups were analyzed using one-way analysis of variance (ANOVA) followed by Tukey’s multiple comparisons post hoc test. Statistical significance is indicated as: *p-value ≤ 0.05, **p-value ≤ 0.01, ***p-value ≤ 0.001 and ****p- value ≤ 0.0001. Unless otherwise stated, experiments were performed with three biological replicates per group. Mouse retinal protein analysis was performed using four biological replicates per group, and Immunohistochemistry analysis was performed using six different images per group.

### Public transcriptomic data analysis

For the primary HREC dataset GSE221861 [48], gene expression data were obtained from the National Center for Biotechnology Information (NCBI) Gene Expression Omnibus (GEO). Differential gene expression was explored using the GEO2R web tool, which compares gene expression between user-defined experimental groups. Samples corresponding to VEGFA-treated and untreated HRECs were selected, and the transcriptional expression of *Ndufa4l2* and *Ndufa4* was examined. The processed expression values generated by GEO2R were downloaded and replotted using GraphPad Prism version 10.

For the mouse P17 OIR RNA-sequencing dataset GSE234447 [36], processed RNA-seq expression data were downloaded from the GEO database. The dataset consisted of retinal samples collected from C57BL/6J mice subjected to the P17 OIR model. Expression levels of *Ndufa4l2* and *Ndufa4* were extracted from the processed dataset and visualized using GraphPad Prism version 10.

To determine the cellular distribution of *Ndufa4l2* expression in the adult mouse retina, the single-cell RNA sequencing dataset GSE243413 [41] was analyzed using the Broad Institute Single Cell Portal. Published visualization tools available through the portal, including Uniform Manifold Approximation and Projection (UMAP)-based feature plots and cell-type distribution plots, were used to examine the expression pattern of *Ndufa4l2* across retinal cell populations in adult C57BL/6J mice. Gene expression was queried directly through the portal without additional computational processing.

## Discussion

In this study, we aimed at identifying changes hyperoxia makes to proteins in the retina that may play a role in hyperoxia-mediated downregulation of proliferation and migration of endothelial cells and finally impact vascular development in OIR. Deep protein TMT-labeled sequencing of the retinal proteome of P12 mouse exposed to normoxia or hyperoxia showed downregulation of NDUFA4L2 in the hypovascularized phase I of OIR, a protein that has not yet been linked to any vascular conditions and OIR. In addition to the retina, we also saw downregulation of NDUFA4L2 in cultured HRECs exposed to hyperoxia in comparison to normoxic controls. Moreover, our reanalysis of a published RNAseq dataset indicated upregulation of *Ndufa4l2* in phase II of OIR. Since hyperoxia in phase I of OIR leads to hypovascularization and returning animals to room air from postnatal day 12 to day 17 in phase II causes excessive vascularization, our findings that NDUFA4L2 is downregulated in phase I and upregulated in phase II correlate with the vascular changes seen in the respective phases. NDUFA4L2 localization to retinal endothelial cells was confirmed by reanalyzing publicly available retinal single cell data set (GSE243413 [41]), further supporting its potential role in endothelial cells. Notably, NDUFA4L2 was detected only in vertebrate species possessing a circulatory vascular system, whereas no orthologs were identified in the invertebrate species analyzed (Supplemental Figure S5a and b). Together with our experimental findings in endothelial cells, this evolutionary conservation supports the notion that NDUFA4L2 has evolved to fulfill specialized functions within the vertebrate vasculature. NDUFA4L2 has a paralog, NDUFA4, with both being subunits of 44 total units in the complex I [58]. Despite having a close sequence similarity with NDUFA4L2, NDUFA4 didn’t show statistically significant changes in the phase I and phase II of OIR. This also further shows that hyperoxia doesn’t simply downregulate the whole complex I but modulates subunits of the complex I to fine-tune its activity.

We then further delved into understanding the mechanistic role NDUFA4L2 plays in vascular development and the pathophysiology of OIR. To do so, we first investigated potential upstream mechanisms that regulate NDUFA4L2 in endothelial cells. As per the majority of published work on NDUFA4L2 in cancer models, NDUFA4L2 is regulated by HIF1α [34, 38–40]. Although HIF1α is downregulated in phase I of OIR, modulating HIF1α with two different pharmacological mechanisms, proteasomal inhibition and prolyl hydroxylases inhibition, did not alter NDUFA4L2 levels in endothelial cells, both in physiological normoxic and pathological hyperoxic conditions. In the OIR model, HIF1α controls VEGFA levels in the retina and both proteins are downregulated in phase I of OIR. This downregulation of VEGFA is considered to be the main cause of vaso-obliteration in phase I of OIR because conceptually, less mitogen means downregulation of VEGF-VEGFR signaling in endothelial cells, and consequential downregulation of VEGFR-driven pathways in endothelial cells. This means an easy fix is to increase VEGFA in phase I of OIR. Interestingly, we and others have shown that VEGFA supplementation alone is not sufficient to rescue the vaso- obliteration phenotype. Notably, the media we used for primary HREC cultures contains VEGFA, and yet hyperoxia still stunted proliferation and migration of ECs in culture, implying the effect of oxygen is downstream of these growth factors. We suspected hyperoxia may also downregulate VEGFA perception by downregulating VEGFR2. Our immunocytochemistry data showed downregulation of VEGFR2. This made us think that the VEGF-VEGFR2 pathway is causative to NDUFA4L2 downregulation. We used a very potent VEGFR2/tyrosine kinase inhibitor, Apatinib, to inhibit VEGFR2 and found statistically significant downregulation of NDUFA4L2 with Apatinib treatment. This implied that VEGFR2 is upstream of NDUFA4L2. We then conducted VEGFA gradient and VEGFR2 siRNA experiments, with both showing no effect whatsoever on NDUFA4L2 levels, indicating that VEGF-VEGFR2 are potentially not upstream of NDUFA4L2. Since Apatinib has also been shown to downregulate PI3K/AKT/mTOR independent of VEGFR2 [59], we suspect that NDUFA4L2 may be regulated by this PI3K/AKT/mTOR signaling. Another possibility is that Apatinib binds a previously unknown receptor that controls NDUFA4L2 levels in a PI3K/AKT/mTOR-dependent manner. To further understand the regulation of NDUFA4L2 by hyperoxia, we looked at PTMs on NDUFA4L2 in normoxic and hyperoxic conditions and found 3 unique sites that are PTM-modified in hyperoxic conditions but not in normoxic conditions. Further functional analysis of these PTM sites is necessary to understand their relevance to NDUFA4L2 stability and activity.

We next looked at the effect of NDUFA4L2 on proliferation and migration of endothelial cells, two main phenotypes seen in OIR. Although NDUFA4L2 has been shown to reverse proliferation defects in cancer models [34, 38–40], our findings show that this is not the physiological role of NDUFA4L2 in endothelial cells. NDUFA4L2 overexpression in primary HRECs rescued migration defects caused by hyperoxic exposure. To understand the mechanistic underpinning of how NDUFA4L2 rescues migration defects, we measured metabolomic changes in primary HRECs overexpressing NDUFA4L2 and compared them with mCherry overexpression controls. NDUFA4L2 is a mitochondrial protein previously shown to control glycolysis in cancer cells [60]. Interestingly, in physiological and pathological hyperoxic conditions, NDUFA4L2 increased the lactate/pyruvate ratio in sub-confluent cultured HRECs, implying an increase in aerobic glycolysis in the sub-confluent proliferative phase. Notably, hyperoxia alone did not significantly alter the lactate/pyruvate ratio or citrate levels in sub-confluent HRECs, consistent with our previous metabolomic analysis of sub-confluent hyperoxia-exposed primary HRECs [61]. The reproducibility of these findings across two independent studies strengthens the conclusion that hyperoxia does not substantially perturb these metabolic parameters in proliferating sub-confluent HRECs. LC-MS and GC-MS analysis of cells in a confluent state further revealed that hyperoxia downregulates IDH flux and NDUFA4L2 rescues IDH flux. Remarkably, these metabolic effects were absent in sub-confluent HRECs —a proliferative state—indicating that αKG and downstream TCA flux regulation by NDUFA4L2 is a metabolic mechanism that rescues migration phenotype only in confluent HRECs. Interestingly, exogenous dimethyl- αKG has been shown to enhance HUVEC proliferation, migration, and tube formation through PI3K/Akt/HIF1α signaling pathway [62], and an αKG-dependent P4HA1-TET2-FBP1 pathway has been shown to regulate the balance between oxidative and glycolytic metabolism during hypoxic angiogenesis [63]. Exogenous αKG also rescues angiogenic dysfunction in endothelial progenitor cells in diabetic mice [64]. As the mechanism we identify, TCA flux at the IDH step, is a distinct and potentially targetable point, supplementing αKG directly, or targeting its upstream kinase once identified, may represent a therapeutic strategy for preventing phase I vaso-obliteration before it progresses to the neovascular phase II, where current anti-VEGF treatments carry their own risks to vascular development. The main findings in this manuscript are depicted in schematic Figure 7.

**Figure 7.**
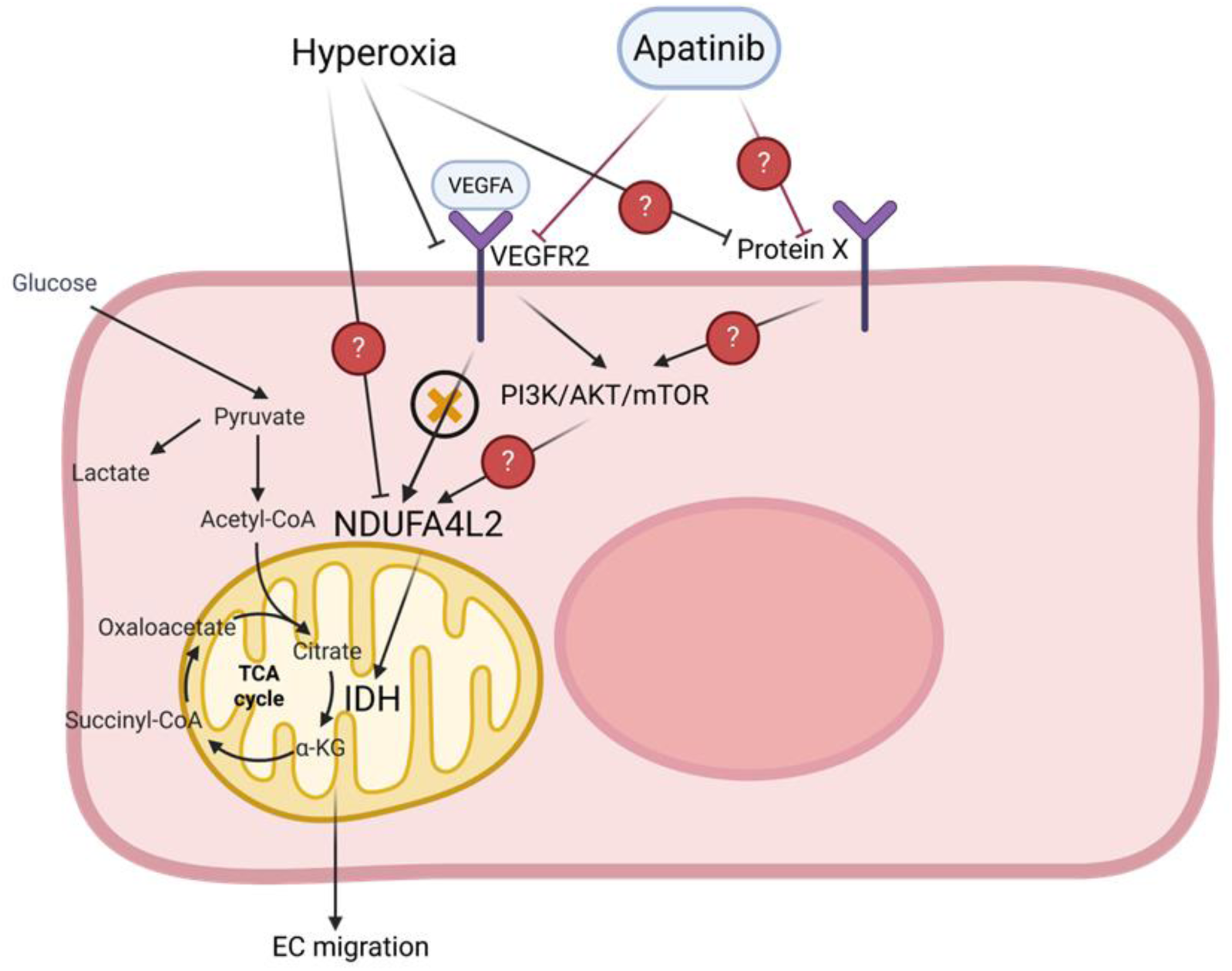
Schematic depicting the potential pathways that control NDUFA4L2 in hyperoxic endothelial cells. The schematic also shows how NDUFA4L2 regulates metabolic changes in IDH flux to rescue migration defects in retinal endothelial cells in OIR. Our findings indicate that hyperoxia downregulates NDUFA4L2 in a VEGF-VEGFR2- independent but Apatinib-dependent manner. Hyperoxia significantly reduced VEGFR2 expression. However, NDUFA4L2 protein levels were not directly regulated by VEGF-VEGFR2 signaling axis as implied by VEGF gradient and siVEGFR2 experiments. In contrast, treatment with Apatinib, a VEGFR2/tyrosine kinase inhibitor, significantly decreased NDUFA4L2 protein levels. Given that Apatinib has been shown to inhibit the PI3K/AKT/mTOR signaling pathway independent of VEGF-VEGFR2 signaling axis, these findings raise the possibility that hyperoxia and Apatinib may regulate NDUFA4L2 expression through an unidentified upstream receptor (Protein X) and potentially via downstream PI3K/AKT/mTOR signaling. Alternatively, hyperoxia may directly reduce NDUFA4L2 protein stability through PTMs, consistent with our PTM analysis. Our findings demonstrate that NDUFA4L2 rescues endothelial cell migration defects by restoring IDH flux and αKG production in retinal endothelial cells, highlighting NDUFA4L2 dependent IDH flux regulation as a potential therapeutic target for preventing Phase I OIR.

## Acknowledgement

This research was supported by National Eye Institute/National Institute of Health grant number R21EY033046 and R01EY035701 to CS, and by Massachusetts Lions Eye Research Fund, Inc. grants to CS.

pLJM1-Empty was a gift from Joshua Mendell (Addgene plasmid # 91980 ; http://n2t.net/addgene:91980 ; RRID:Addgene_91980)

psPAX2 was a gift from Didier Trono (Addgene plasmid # 12260 ; http://n2t.net/addgene:12260 ; RRID:Addgene_12260)

pCMV-VSV-G was a gift from Bob Weinberg (Addgene plasmid # 8454 ; http://n2t.net/addgene:8454 ; RRID:Addgene_8454)

Some figures in this manuscript were created using BioRender.com

---

**Supplemental Figure S1.**
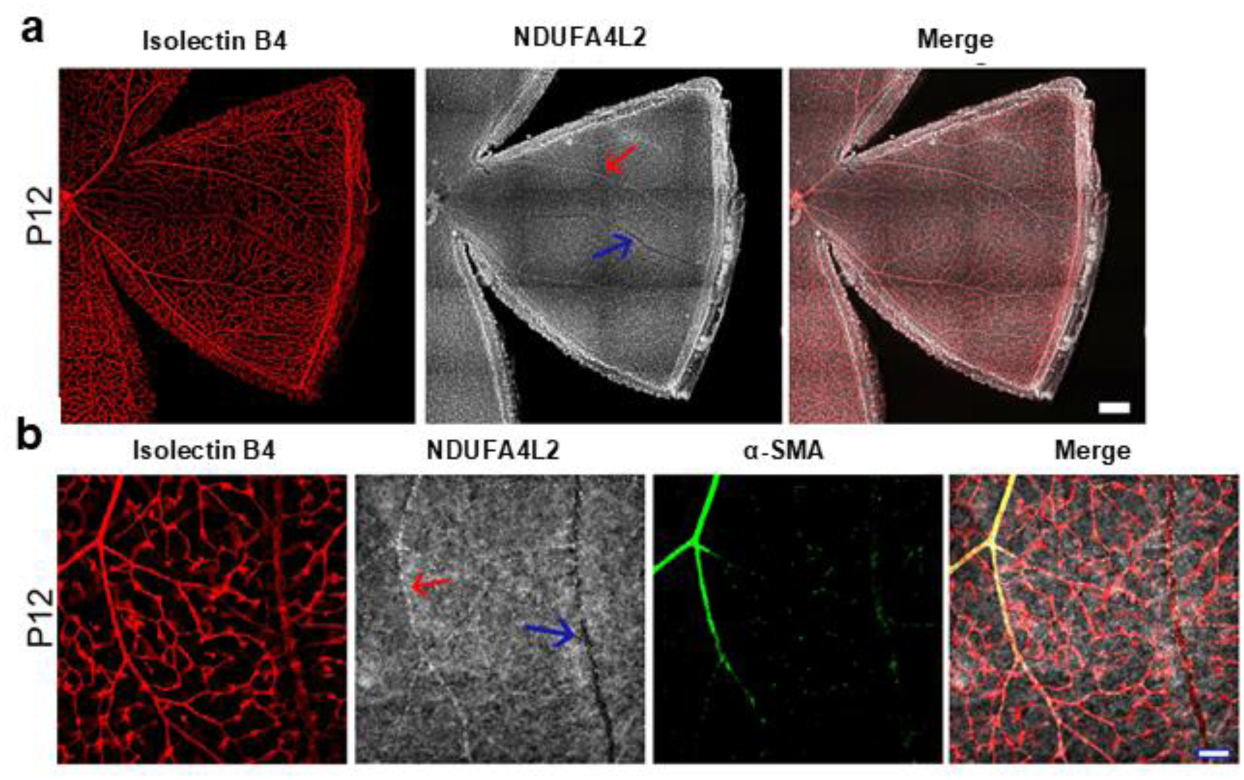
NDUFA4L2 localizes in the artery of the P12 mouse retina. a) Representative P12 retinal flatmounts of C57BL6J (n=6) mice immunostained for isolectin b4 and NDUFA4L2, overlayed image showing NDUFA4L2 vascular expression pattern. Arrows indicate arteries (in red) and veins (in blue). b) Representative P12 retinal flatmounts of C57BL6J (n=6) mice immunostained for isolectin b4, NDUFA4L2, and alpha-smooth muscle actin (αSMA), showing NDUFA4L2 vascular expression pattern in arteries (red arrow) versus veins (blue arrow). Scale bar: A is 200 µm and B is 75 µm.

**Supplemental Figure S2.**
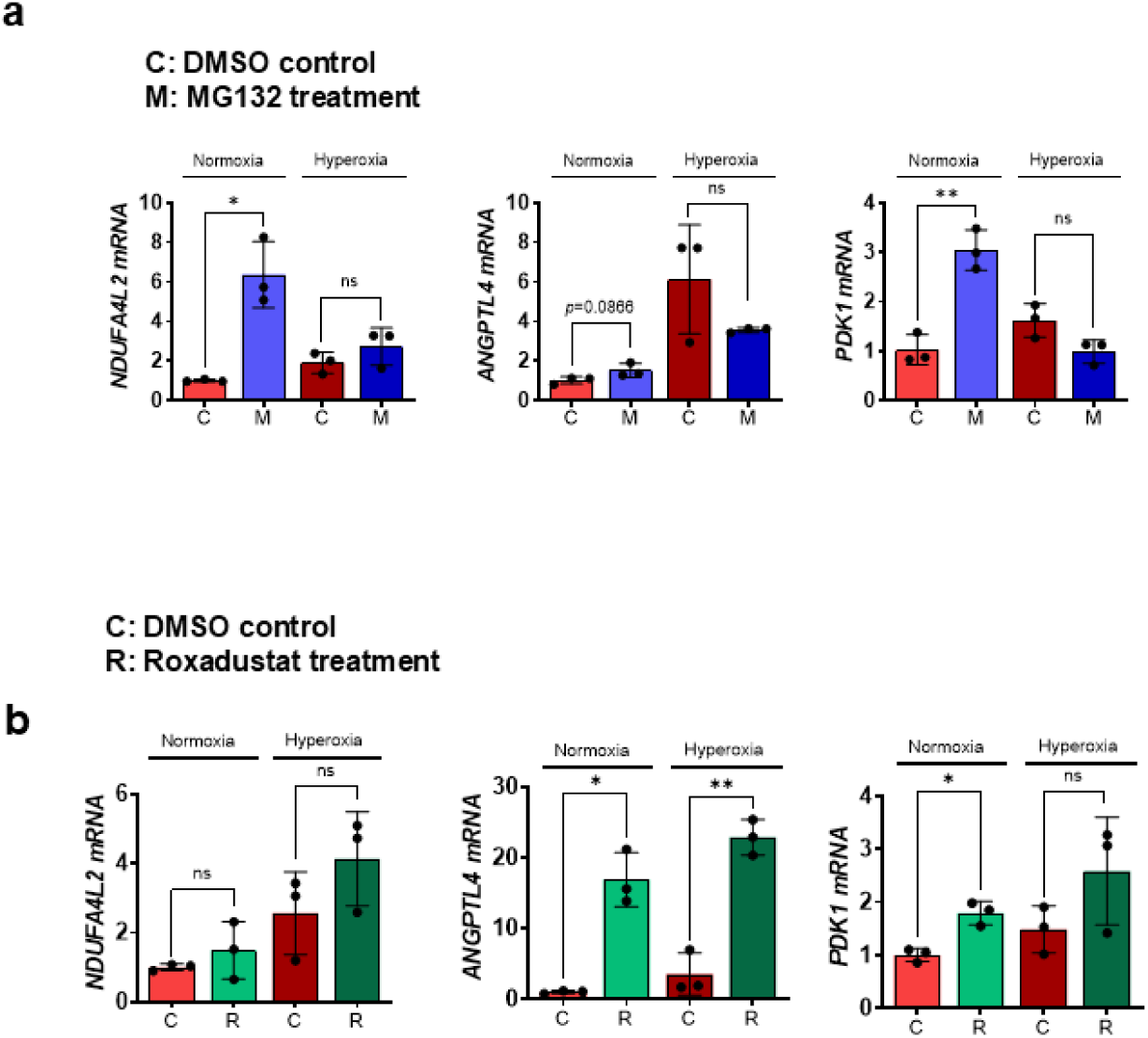
MG132 and Roxadustat induced the transcription levels of HIF1α target genes, indicating active HIF1α in these conditions. a) Quantitative RT-PCR (qRT-PCR) result showed that the low dose of MG132 treatment in primary HRECs increased the transcription levels of *NDUFA4L2* in normoxia and some HIF1α downstream targets. b) qRT-PCR of HIF1α downstream targets showed higher expression in response to Roxadustat treatment. Data are presented as mean ± SD (n = 3 per group). Statistical significance was determined using an unpaired two-tailed Welch’s t-test. Statistical significance is indicated as: *p-value ≤ 0.05, and **p-value ≤ 0.01. NS indicates no significant difference. ANGPTL4, Angiopoietin-like 4; PDK1, Pyruvate dehydrogenase kinase 1

**Supplemental Figure S3.**
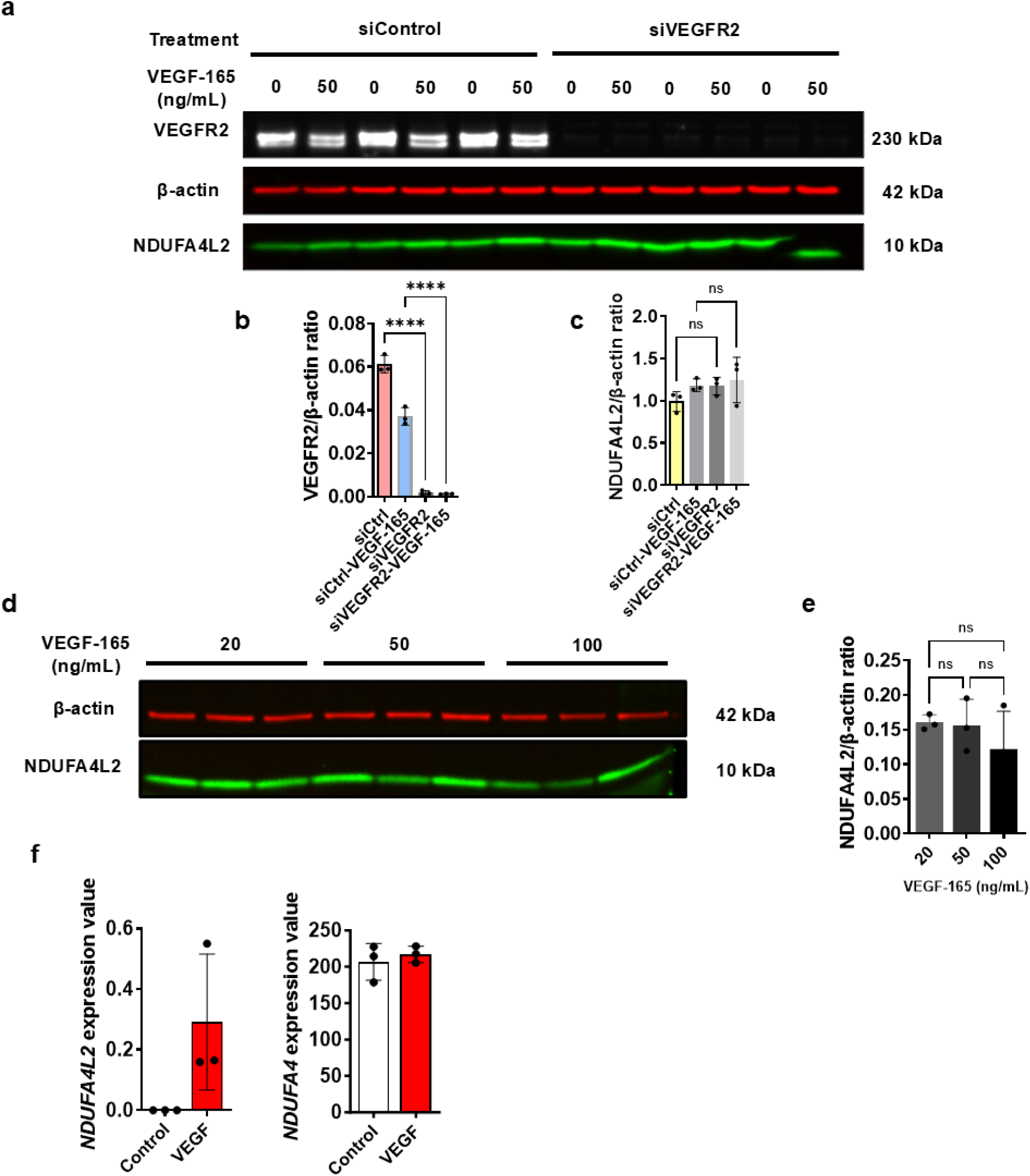
We didn’t find NDUFA4L2 protein regulation by VEGF-VEGFR2 signaling in our analysis, but its transcription levels were changed by VEGF dependent manner. a) Knockdown of VEGFR2 in primary HRECs showed very strong downregulation of VEGFR2 protein but did not change NDUFA4L2 protein levels, regardless of human VEGF-165 treatment. b) quantification of VEGFR2 levels from siVEGFR2 western blot showing strong downregulation of VEGFR2 protein. c) quantification of NDUFA4L2 levels from siVEGFR2 western blot showing no statistically significant differences. d) Supplementing media with different concentrations of VEGF-165 did not alter NDUFA4L2 protein levels. e) quantification of NDUFA4L2 levels from VEGF-165-treated primary HRECs showing no statistically significant differences. Data are presented as mean ± SD (n = 3 per group). Comparisons among four groups were analyzed using one-way ANOVA followed by Tukey’s multiple comparisons post hoc test. Statistical significance is indicated as: ****p- value ≤ 0.0001. NS indicates no significant difference. f) Re-analyzed data from GEO dataset GSE221861 by Rameshekar et al.[48] showed the induction of *NDUFA4L2* transcription level in primary HRECs in response to VEGFA, but not its paralog *NDUFA4,* indicating that this may be regulated at transcriptional levels but not at protein levels. Data are presented as mean ± SD (n = 3 per group).

**Supplemental Figure S4.**
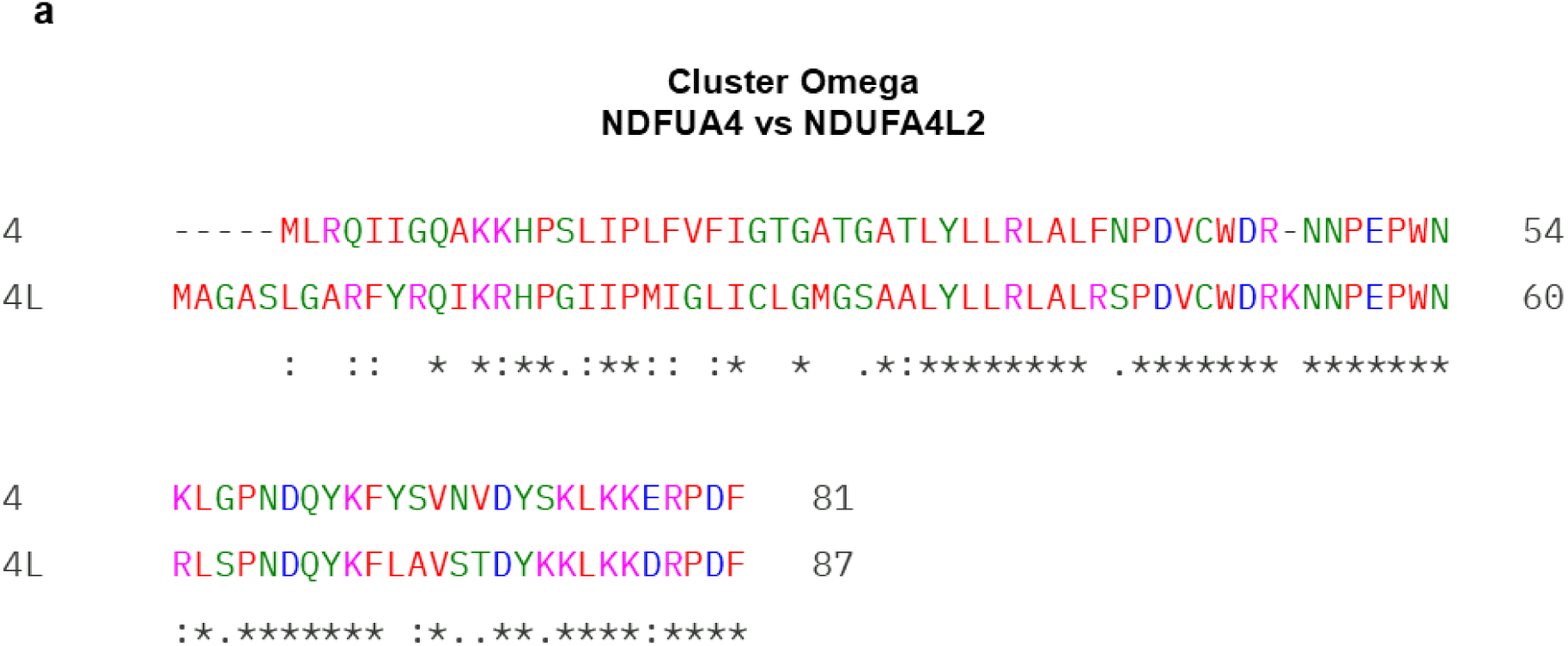
NDUFA4L2 has some unique amino acids in its sequence compared to its paralog, NDUFA4, and these may be regulatory. a) Protein sequences of NDUFA4L2 and NDUFA4 from Uniprot and aligned them in Clustal Omega, showing some regions of dissimilarity between the two

**Supplemental Figure S5.**
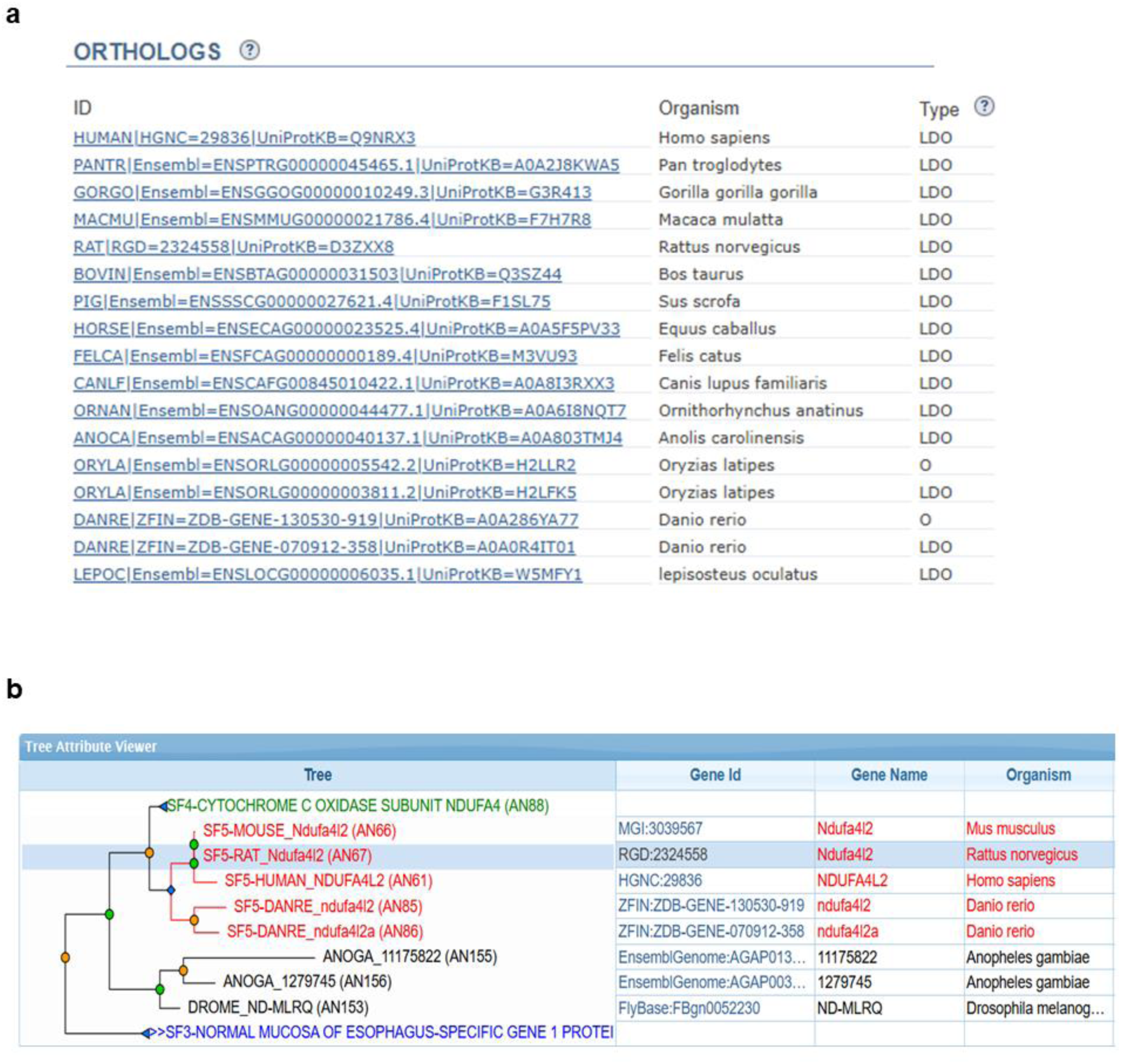
NDUFA4L2 is a vertebrate-specific paralog restricted to organisms bearing vasculature. a) The ortholog table generated using the PANTHER database shows species in which an NDUFA4L2 ortholog is present. Least diverged orthologs (LDO) and orthologs (O) were identified across mammals, a monotreme, a reptile, and multiple fish species, all of which possess a circulatory vascular system of any kind. b) Gene tree from the PANTHER database shows the evolutionary relationship between NDUFA4L2 (subfamily SF5) and its paralog NDUFA4 (subfamily SF4). NDUFA4L2 orthologs form a distinct vertebrate-specific clade (red), separate from the ancestral ND-MLRQ gene found in invertebrate species lacking a closed vasculature (*Anopheles gambiae*, *Drosophila melanogaster*). Gene ID, gene name, and organism are shown for each node.

**Supplemental Figure S6.**
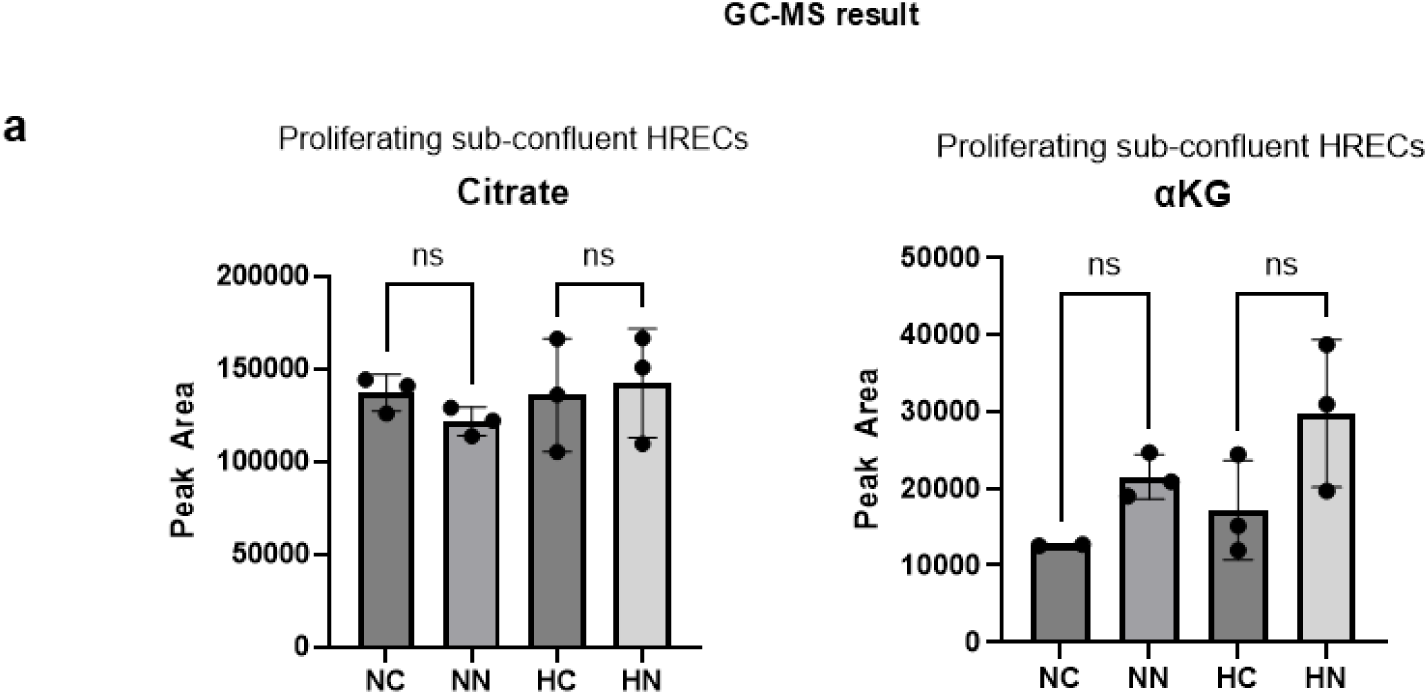
IDH flux is not impacted in proliferating sub-confluent HRECs and shows no significant changes in response to NDUFA4L2 overexpression. a) GC-MS result indicates that NDUFA4L2 did not restore citrate and αKG in proliferating sub-confluent HRECs. Data are presented as mean ± SD (n = 3 per group). Comparisons among four groups were analyzed using one-way ANOVA followed by Tukey’s multiple comparisons post hoc test. NS indicates no significant difference. Legends: NC, normoxia mCherry OE; NN, normoxia NDUFA4L2 OE; HC, hyperoxia mCherry OE; HN, hyperoxia NDUFA4L2 OE.

